# Multiple mechanisms contribute to isolation-by-environment in the redheaded pine sawfly, *Neodiprion lecontei*

**DOI:** 10.1101/2023.05.22.541587

**Authors:** Robin K. Bagley, Melanie N. Hurst, Jeremy Frederick, Jordan Wolfe, John W. Terbot, Christopher J. Frost, Catherine R. Linnen

## Abstract

Isolation by environment (IBE) is a population genomic pattern that arises when ecological barriers reduce gene flow between populations. Although current evidence suggests IBE is common in nature, few studies have evaluated the underlying mechanisms that generate IBE patterns. In this study, we evaluate five proposed mechanisms of IBE (natural selection against immigrants, sexual selection against immigrants, selection against hybrids, biased dispersal, environment-based phenological differences) that may give rise to host-associated differentiation within a sympatric population of the redheaded pine sawfly, *Neodiprion lecontei*, a species for which IBE has previously been detected. We first characterize the three pine species used by *N. lecontei* at the site, finding morphological and chemical differences among the hosts that could generate divergent selection on sawfly host-use traits. Next, using morphometrics and ddRAD sequencing, we detect modest phenotypic and genetic differentiation among sawflies originating from different pines that is consistent with recent, *in situ* divergence. Finally, via a series of laboratory assays – including assessments of larval performance on different hosts, adult mate and host preferences, hybrid fitness, and adult eclosion timing – we find evidence that multiple mechanisms contribute to IBE in *N. lecontei*. Overall, our results suggest IBE can emerge quickly, possibly due to multiple mechanisms acting in concert to reduce migration between different environments.

## Introduction

Isolation by distance (IBD)—a pattern in which genetic similarity declines as a function of geographic distance between individuals—is one of the most ubiquitous population genetic patterns in nature (Meirmans, 2012; M. A. Peterson & Denno, 1998; Sexton et al., 2014). The primary mechanism generating this pattern is geographically restricted dispersal: as gene flow declines between increasingly distant locations, genetic differentiation can accumulate via drift (Wright, 1943, 1946). Independent of geographical distance, environmental differences can also restrict gene flow between locally adapted populations via multiple mechanisms, such as biased dispersal and natural and sexual selection against immigrants and hybrids. When this occurs, individuals from dissimilar environments will tend to be more genetically differentiated than individuals from similar environments, a pattern called isolation by environment (IBE) (Bradburd et al., 2013; Sexton et al., 2014; Wang & Bradburd, 2014; Wang & Summers, 2010).

Compared to a rich literature documenting both IBD patterns (Battaglia et al., 2008; Jauker et al., 2009; Moore et al., 2008; Svenning et al., 2008; Sydenham et al., 2017) and mechanisms that cause geographically restricted dispersal (Baguette & Van Dyck, 2007; Bowler & Benton, 2005; Clobert et al., 2009; Matthysen, 2013; Pflüger & Balkenhol, 2014; Ronce & Clobert, 2013), research into IBE is more limited. Nevertheless, a growing number of studies are finding evidence of IBE (e.g., Bagley et al. 2017; Prunier et al. 2017; Moncada et al. 2021; Mancilla-Morales et al. 2022), raising the possibility that this pattern is just as ubiquitous—if not more so—than IBD (Sexton et al., 2014; Shafer & Wolf, 2013). Still, major challenges persist when studying IBE. First, geography and ecology are often strongly correlated, making it difficult to disentangle their individual effects. Several approaches have been proposed to account for this spatial autocorrelation [e.g., Mantel and partial Mantel tests (Mantel, 1967; Smouse et al., 1986; Sokal, 1979), BEDASSLE (Bradburd et al., 2013), MMRR (WANG, 2013), SUNDER (Botta et al., 2015)], but their application remains limited. Second, and perhaps more importantly, detection of an IBE pattern does not in itself reveal which of several non-mutually exclusive mechanisms gave rise to the observed pattern. However, few studies have gone beyond documenting IBE to test underlying mechanisms.

Wang and Bradburd (2014) described four potential mechanisms that may generate the IBE pattern via reducing effective migration between divergent environments. First, when populations are locally adapted to a specific environment, individuals from that population may fare poorly when they disperse to an alternative environment (Kawecki and Ebert 2004; Nosil et al. 2005; Zhang et al. 2017; also see Kawecki 1997). Selection against non-locally adapted immigrants will tend to decrease gene flow between dissimilar habitats, with the magnitude of that selection increasing with greater ecological distance between the environments (Crispo et al., 2006; Wang et al., 2013). Second, immigrants may be sexually selected against if they originated from source population experiencing divergent sexual selection. Sexual selection against immigrants can act alongside natural selection, for example if the choosy sex tends to prefer individuals with traits well suited to the local habitat (Ingleby et al., 2010; Jia & Greenfield, 1997; Nosil et al., 2005). Alternatively, sexual signals themselves may be optimized for local environments, thereby reducing signaling efficacy—and reproductive success—when individuals display these signals in different environments (e.g., Boughman 2001; Seehausen et al. 2008; Pires et al. 2019). A third mechanism that can produce an IBE pattern is when natural or sexual selection acts against hybrid offspring produced by parents from different environments. Hybrid individuals may, for example, exhibit an intermediate phenotype unsuitable for either parental habitat, reducing survival or opportunities for attracting mates (e.g., McBride and Singer 2010; Jacquemyn et al. 2018; Chhina et al. 2022). Finally, IBE can be generated if individuals are more likely to disperse to a similar environment than to a different environment, either via genetically based habitat preferences (e.g., Feder and Forbes 2007; Edelaar et al. 2008; Bolnick and Otto 2013) or via plastic responses to the natal habitat (Benard & McCauley, 2008; J. M. Davis & Stamps, 2004; Merrick & Koprowski, 2016).

In addition to the four IBE mechanisms described by Wang and Bradburd (2014), habitat-related differences in phenology can also produce an IBE pattern (Sexton et al., 2014). Such differences can arise via two non-mutually exclusive routes. First, phenological differences among populations in different environments could result from divergent selection and local adaptation. For example, in the apple maggot fly, *Rhagoletis pomonella*, heritable differences in adult eclosion time synchronize host races with the availability of ripe fruit for oviposition, which differs among host plants (Doellman et al., 2018; Feder et al., 1993, 1994). Second, even in the absence of genetic differences in developmental timing, developmental plasticity in response to environmental variables could give rise to differences in reproductive periods between populations living in different habitats. For example, shifts in flowering time may arise in plants occupying niches with differing environmental conditions (e.g., temperature, pH, moisture level) that affect plant physiology (Gavrilets & Vose, 2007; Levin, 2009; Rafferty et al., 2020; Silvertown et al., 2005). Regardless of the source of the phenological shift, when populations differ in the timing of their reproductive periods, gene flow will be reduced between dissimilar environments and a pattern of IBE can be produced (Boumans et al., 2017; Taylor & Friesen, 2017; Y. M. Zhang et al., 2018).

At present, the relative importance of the five different IBE mechanisms remains unknown because very few studies have evaluated IBE mechanisms at all, let alone multiple potential IBE mechanisms in the same system (Wang & Bradburd, 2014). To this end, we evaluate multiple IBE mechanisms in *Neodiprion lecontei,* an experimentally tractable species for which an IBE pattern was previously detected (Bagley et al., 2017). *Neodiprion* (Hymenoptera: Diprionidae) is a Holarctic genus of sawflies that specialize on pines. Like many plant-feeding insects, *Neodiprion* sawflies are closely associated with their host plants throughout their life cycle: adults mate on the host plant, females lay eggs into pockets cut within host needles, and larvae consume the needles during development before spinning cocoons on or beneath the host (Benjamin, 1955; Coppel & Benjamin, 1965; Knerer, 1993; Wilson et al., 1992). Most species also feed on only one or a small handful of host plant species (Linnen & Farrell, 2010; Smith, 1993). Due to this high degree of specialization and intimate, lifelong relationship with their host plants, it has long been hypothesized that host adaptation is a primary driver of population differentiation and speciation in *Neodiprion* sawflies (Ghent and Wallace 1958; Alexander and Bigelow 1960; Knerer and Atwood 1972, 1973; Bush 1975a,b). Consistent with this hypothesis, changes in host use are associated with speciation events in the genus (Linnen & Farrell, 2010), and divergence in host-use traits contributes to both prezygotic isolation (Glover et al., 2023) and extrinsic postzygotic isolation(Bendall et al., 2017).

Although *Neodiprion lecontei* is a pine generalist compared to other species in the genus (Linnen & Farrell, 2010; Wilson et al., 1992), *N. lecontei* populations collected from different pine species tend to be more genetically dissimilar than those collected from the same pine species, after controlling for historical isolation and geographic distance (Bagley et al., 2017). To understand why this pattern exists, we use field observations and laboratory experiments to evaluate potential IBE mechanisms in *N. lecontei*. To remove the effect of geography entirely, we focus on a single location where *N. lecontei* was observed feeding on three different *Pinus* species. To evaluate potential host-related sources of divergent selection, we first characterize the morphology, volatile chemistry, and resin content of the three *Pinus* hosts. To characterize population structure and evaluate IBE at this single site, we generate genome-wide genetic data via double-digest restriction-associated DNA (ddRAD) sequencing. Finally, to understand mechanisms that give rise to host-associated differentiation (a specific type of IBE) among *N. lecontei* collected on three pine species, we evaluate: (1) host-based performance differences (natural selection against immigrants), (2) mate preferences (sexual selection against immigrants), (3) hybrid survival (natural selection against hybrids), (4) female host preferences (habitat-based dispersal bias), and (5) adult eclosion patterns (habitat-related differences in phenology). Taken together, our results suggest that multiple IBE mechanisms can and do act in concert, possibly facilitating its rapid emergence in natural populations.

## Materials and Methods

### Study site and host trees

Our study site was located at the at the University of Kentucky’s Arboretum and State Botanical Gardens, established in 1991 in Lexington, KY. Spanning a transect of ∼130m, the “Trail of Pines” (38.0167°N, 84.5047°W) has three pine species, all of which are native to Kentucky: *Pinus echinata* (shortleaf pine), *P. virginiana* (Virginia pine), and *P. rigida* (pitch pine). The trees were planted in the mid-to-late 1990s (T. Rounsaville, personal communication) near each other, with the branches of some trees physically intermingling [Supplemental Figure 1]. The primary study period was between 2012 and 2015, during which *N. lecontei* larvae were abundant on all three species. Although we do not know exactly when sawflies first colonized the study site, given the age of the trees, the age of the Trail of Pines population of *N. lecontei* was likely no more than ∼10-15 years old (and possibly much younger) at the start of the study. With 2-3 generations per year, on average, in Lexington (RKB and CRL, personal observation), the maximum age of the population is 30-45 generations.

To gain insight into potential host-related selection pressures that could give rise to IBE among sawflies feeding on the three different pine species, we described morphological and chemical differences between the pine species. First, because needle width affects oviposition success of *N. lecontei* females (Bendall et al., 2017; Glover et al., 2023), we asked whether needle width differed among the three pine species. From each individual pine tree in the Trail of Pines (*P. echinata N* = 4, *P. rigida N* = 2, *P. virginiana N* = 3), we measured the width of 10 needles with digital calipers (Mitutoyo CD-6”PMX). Because we used greenhouse-grown seedlings of these same species in our assays (see below), we also measured the width of seedling needles so that we could make comparisons between trees at our collection site (potential sources of divergent selection on wild-caught sawflies) and the seedlings used in our assays. For each of the three species of pine, we measured 10 needles from each of 10 randomly selected seedlings.

We analyzed the needle data – and, unless otherwise noted, all other data – in R version 4.2.1 (R Core Team, 2022). To determine how needle width varied as a function of life stage (mature vs. seedling) and pine species, we used the *lmer* function (lmerTest v 3.1-3; Kuznetsova et al. 2017) to fit a mixed-effect model to the needle width measures, with individual tree as a random effect and pine species, life stage, and their interaction as fixed effects. We used a Type III Analysis of Variance (ANOVA) to assess the significance of the fixed effects, followed by the *emmeans* function (emmeans v1.8.0; Lenth 2020) for *post hoc* pairwise comparisons among: (a) pairs of pine species, (b) life stages within pine species, and (c) pairs of pine species within life stages. All *post hoc* comparisons used Benjamini-Hochberg correction for multiple testing.

Next, to determine whether volatile profiles differed among hosts, which could facilitate divergence in female host preferences, we collected the volatiles emitted from trees planted along the path of pines (*P. echinata*, *n* = 4; *P. rigida*, *n* = 4; and *P. virginiana, n* = 4). We conducted headspace sampling by enclosing a set of needles in a caprolactum bag and loosely securing the open end. Each bag had a volatile collecting filter (containing ∼30mg PorapakQ) secured to the bag, and air from the headspace was pulled through the collection filter using small diaphragm vacuum pumps (Karlsson Robotics) powered by 5A, 12V batteries that generated approximately 3 L/min of flow (Frost, 2023). Volatile collections lasted 15 min each. Filters were extracted with 150 μl dichloromethane containing 10ng/μl of nonyl acetate as an internal standard (Frost et al., 2007). Volatile analytes were resolved and detected using a 7890B gas chromatograph (GC) coupled to a 5977A single-quadrupole mass spectrometer (MS) (Agilent Technologies). The GC operated in splitless mode with an inlet temperature of 250°C and a DB-5 column (30 m length, 0.25 mm diameter) with a flow of 1.0 ml/min, and temperature programming initiated at 35°C for 1 min, ramped 20°C/min to 250°C, with a 5 min hold at 250°C. The MS EI ion source (70 eV) and quadrupole temperatures set at 230°C and 150°C, respectively. Mass spectra were collected in scanning ion mode (*m/z* 40 and 500) and deconvoluted using MassHunter Quantitative Analysis software (vB.07.00, Agilent Technologies). Preliminary compound identities were determined from the NIST14 database with spectral match thresholds for metabolites at >85% (Frost et al., 2012). We compared the total volatile profiles using non-metric multi-dimensional scaling (NMDS), a robust ordination method that has been commonly applied to the multivariate analysis of volatile profiles (Bricchi et al., 2010; Minchin, 1987). NMDS was performed using the *metaMDS* function (implemented in the R package vegan v2.6-4; Oksanen 2010) with 95% confidence ranges generated using *veganCovEllipse* (see DRYAD for R code). For analysis of individual volatiles, we fit a linear model (*lm*) for each compound with tree species as the fixed factor. We used a Type II ANOVA (as implemented in the R package car v3.1-1; Fox and Weisberg 2019) to assess the main effect across the three pine species, followed by *post hoc* pairwise comparisons (emmeans v1.8.1-1).

Finally, to determine whether resin content differed among the three pine species, which could differentially impact larval growth and survival (Larsson et al., 1986), we quantified resin content from branch clippings collected from exemplars of each species. These clippings were collected as a part of a broader survey of pine traits in eastern North America (Glover et al., 2023), but we focus specifically here on samples taken from *P. echniata*, *P. virginiana,* and *P. rigida*. For each tree species, we sampled 10 clippings each from 3-4 geographically widespread locations (Supplemental Table S1). We sampled each site in May (5/8/17 - 5/21/17) and August (8/3/17 - 8/15/17), both months during which *N. lecontei* can be found in the field (Wilson et al., 1992). After collection, clippings were placed into individual plastic bags and stored on ice until taken to the lab. Upon return to the lab, clippings were stored at 4°C until resin data was collected.

We quantified total non-volatile resin content using methods adapted from Moreira (2012, 2014). For each sample, we first weighed approximately 3g of pine needles and finely cut them using a clean razor blade. The needle mass was then placed into a test tube and 6mL of 95%-99% purity hexane added. The extraction was then placed into an ultrasonic bath for 15 minutes at room temperature. Samples were then left capped in a fume hood for 24 hours. We then filtered hexane extract into a second test tube through a GF/D filter and the vegetative material returned to the original test tube. Hexane (6mL) was once again added to the vegetative material and the test tubes placed into an ultrasonic bath for ten minutes. This process was repeated once more for a total of three hexane extracts per sample. Hexane extracts were placed in a fume hood covered with a delicate task wipe to prevent contamination and left until all hexane had evaporated, which was approximately 5 days. Across hundreds of samples that were processed, slight deviations from this protocol occasionally occurred and are noted in Supplemental Table S1.

To estimate the total non-volatile resin content per sample, we weighed the three dried extracts and took their sum. After the final filtration, the tube containing the vegetative material was dried at 40°C until weight stabilized. We then weighed these samples and subtracted the weight of empty tubes to determine the dry weight of the vegetative mass. The percentage resin content was calculated as: (total non-volatile resin content/dry weight of the vegetative mass) * 100. After excluding samples with missing data, the final sample size for each host/month combination ranged between *n* = 29 and *n =* 40.

To determine whether resin content differed among the three host species, we used the *lm* function in R to fit a linear model to the resin content data, with host species, month, and their interaction as predictors. Because model residuals were not normally distributed for untransformed data, we fit a linear model to square-root transformed resin data. To evaluate the significance of model terms, we used Type III ANOVA (car package). We then used the emmeans package for *post hoc* comparisons that evaluated whether there were differences in resin content among pairs of host species either (1) without accounting for sampling period or (2) within each of the two sampling periods, with the Benjamini-Hochberg method for adjusting p-values.

### Sawfly sampling and propagation

Sawfly colonies (i.e., distinct clusters of feeding larvae) were collected from *P. echinata*, *P. rigida*, and *P. virginiana* at the field site between 2012 and 2015 as early-to-late instar feeding larvae (Supplemental Table S2). A subset of larvae from some colonies was preserved in 100% ethanol for population genetic analysis. The remaining larvae were returned to the lab and reared in plastic boxes (32.4 cm × 17.8 cm × 15.2 cm) with mesh lids and provided clippings of their source host species *ab libitum*. Cocoons were collected three times weekly and stored in individual gelatin capsules until emergence. Larvae and cocoons were kept in walk-in environmental chambers maintained at 22**°**C, and an 18:6 light-dark cycle. Cocoons were checked daily for emergence, and live adults (which are non-feeding) were stored at 4**°**C to prolong life until needed for propagation or experimental assays.

For IBE mechanism assays (see below; Figure 1), we established lab lines from larval colonies that were originally collected from each host species (hereafter, referred to by source host as “Shortleaf”, “Pitch”, and “Virginia” lines) and reared them for an additional one to two generations on a common non-natal host in the lab. Briefly, each host line was produced by releasing male and female adults reared from multiple larval colonies (to maximize genetic diversity) into mesh cages containing multiple seedlings of a host that does not occur in Kentucky, *P. banksiana* (jack pine). The adults were allowed to mate and oviposit freely. Upon hatching, larvae from these cages were transferred into plastic boxes and reared as described above on clippings of field-collected *P. banksiana*. We reared all colonies on the same host species to control for the impact of rearing host on host-related phenotypes. We chose *P. banksiana* as the shared host because it is a primary host for *N. lecontei* (Wilson et al., 1992), is a suitable host for most *Neodiprion* species (Knerer, 1984), and because seedlings of this host could be purchased year-round.

**Figure 1.**
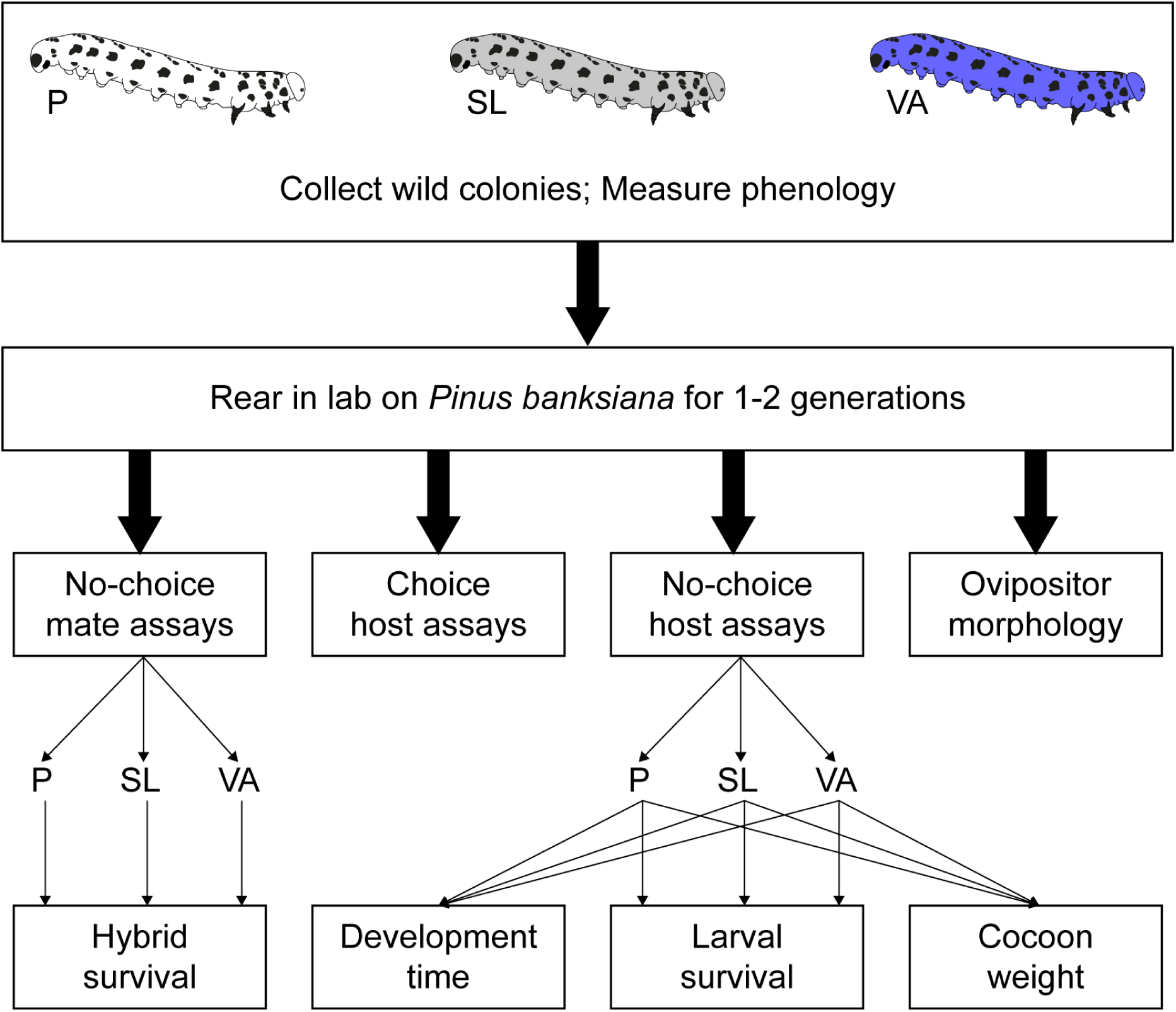
Overview of IBE mechanism assays. All individuals used in our assays were collected at the “Trail of Pines” field site on *Pinus rigida* (pitch pine = “Pitch” line = P, shown in white throughout manuscript), *P. echinata* (shortleaf pine = “Shortleaf” line = SL, shown in grey throughout manuscript), or *P. virginiana* (Virginia pine = “Virginia” line = VA shown in blue throughout manuscript), and then reared as described in the text in the laboratory for 1-2 generations on a common, non-local host (*P. banksiana*, jack pine) before being used in various assays. Arrows are drawn to indicate the flow of individuals and their offspring into and between assays.

### Assessment of Population Structure and Genomic Differentiation

#### DNA extraction, library preparation, and genotyping

A large dataset of SNP markers was prepared using the same extraction and double-digest RAD (ddRAD) sequencing approaches described in Lindstedt et al. (2022) and Bendall et al. (2022). Briefly, 58 Arboretum individuals prepared following a modified version of the original ddRAD protocol (B. K. Peterson et al., 2012), and labeled with one of 48 unique, variable-length (Burford Reiskind et al., 2016) in-line barcodes during adapter ligation (Supplemental Tables S3, S4). Barcoded libraries were pooled, size selected (average fragment size = 379 +/− 76bp) and amplified using multiplex read indices (Supplemental Tables S3, S5). We also included a string of 4 degenerate bases next to the Illumina read index to allow for the detection of PCR duplicates (Schweyen et al., 2014). Two lanes of 150bp single-end reads from an Illumina HiSeq 4000 were obtained for the libraries at the High-Throughput Sequencing and Genotyping Unit at the University of Illinois.

Raw sequence reads were quality filtered and trimmed using the *process_radtags* module in STACKS (v1.46; Catchen et al. 2013). Surviving reads were then aligned to a high-coverage, scaffolded genome assembly for *N. lecontei* (Vertacnik et al. 2016; Linnen et al. 2018; coverage: 112x; scaffold N50: 244kb; GenBank assembly accession: GCA_001263575.1) using BOWTIE2 (v2.3.1; Langmead and Salzberg 2012). Uniquely-mapping, high quality reads (MAPQ ≥30) were extracted with SAMTOOLS (v1.3; Li et al. 2009), and putative PCR duplicates removed. RAD loci were then constructed using the *ref_map.pl* STACKS pipeline (v1.46; Catchen et al. 2013).

After an initial round of SNP calling, we evaluated ploidy and missing data levels using VCFtools (v0.1.14b; Danecek et al. 2011) and excluded seven individuals missing data at >60% of SNP loci and two putative haploid individuals (Supplemental Table S3). Our final RAD dataset consisted of 49 individuals (15 Shortleaf, 13 Pitch, and 21 Virginia). We applied several additional filters to these individuals, excluding all sites missing data in 30% or more of individuals, all sites with a minor allele frequency less than 0.05, and all sites violating Hardy-Weinberg equilibrium for heterozygote excess significant at the 0.01 level. Finally, to minimize linkage disequilibrium between SNPs, we included only one randomly selected SNP per RAD locus. Data processing and all other bioinformatic analyses were performed on the University of Kentucky’s Lipscomb High Performance Computing Cluster.

#### Discrete Population Structure Analyses

To evaluate the possibility that our study population was seeded by host-specialized populations that diverged elsewhere and were already genetically distinct, we asked whether there was any evidence of discrete population structure. We used the maximum-likelihood-based clustering algorithm implemented in the program ADMIXTURE (v1.3.0; Alexander et al. 2009) to determine the proportion of ancestry for each individual from *K* ancestral populations without *a priori* designation. We performed 100 independent runs for values of *K* from 1 through 10. The optimal *K* was selected as described in the ADMIXTURE manual, by comparing the 5-fold cross-validation (CV) error across different values of *K*. To determine assignment stability and visualize primary and secondary solutions across the 100 replicates of each *K*, we used the main pipeline of CLUMPAK (v1.1; Kopelman et al. 2015).

#### Isolation by Environment (IBE): host-associated genetic differentiation

Having found no evidence of discrete population structure indicative of previous isolation among the different host lines (see results), we next asked whether there was any evidence of host-associated differentiation, a specific type of IBE. To do so, we used custom R scripts (available on DRYAD) to compute the Hudson estimator of F_ST_ (Hudson et al., 1992) for each pair of sawfly lines, following Bhatia et al. 2013. To evaluate the significance of observed F_ST_ estimates, we permuted individuals among populations, recalculating F_ST_ in each permutation. P-values were calculated from the proportion of 10,000 permutations that had F_ST_ values greater than or equal to the observed F_ST_ values.

A pattern of isolation by environment is expected to emerge when reduced effective migration between different environments enables genetic differentiation to accumulate via drift and selection. When divergent selection contributes to IBE (e.g., via immigrant or hybrid inviability), differentiation at selected loci should be elevated relative to the rest of the genome (Wang & Bradburd, 2014). To divide the genome into putatively selected and putatively neutral regions, we filtered our RAD markers based on their proximity to annotated genes in the *N. lecontei* genome. We considered genic SNPs (putatively selected) to be within 1-kb of an annotated gene and “intergenic” SNPs to be >5kb away from an annotated gene. We chose these windows based on a median gene length of ∼3.6kb in the NCBI *Neodiprion lecontei* Annotation Release 100 and because the window sizes represented the greatest stringency we could manage without losing unacceptable amounts of data. To obtain the locations of genes in the *N. lecontei* genome, we downloaded a version of the NCBI *Neodiprion lecontei* Annotation Release 100 that had been updated to the scaffolded assembly (GCA_001263575.2), available on NAL@i5k. We then used the GFF2bed script in BEDOPs v 2.4.35 to convert the gff annotation file to a bed file, which we modified in Microsoft Excel to include windows corresponding to our definition of “genic” regions (1000 base pairs subtracted from the start position of the gene and added to the end) and “intergenic” regions (5000 base pairs subtracted and added to the start and end, respectively). We then used these bed files and the include-bed and exclude-bed commands in VCFtools to filter our RAD SNPs, resulting in 4,588 genic SNPs and 3,262 intergenic SNPs.

After generating genic and intergenic datasets, we computed Hudson’s F_ST_ and evaluated significance between each pair of sawfly lines as described above. Finally, to determine whether per-locus estimates of F_ST_ between a particular pair of hosts were elevated in genic regions compared to intergenic regions, we first used the *--weir-fst-pop* command in VCFtools to calculate per-locus F_ST_ for each pair of populations. Then, we used another custom R script (available on DRYAD) to permute F_ST_ values across loci. We then calculated the p-value (one-tailed) as the proportion of 10,000 permutations for which the difference between the average genic F_ST_ and average intergenic F_ST_ was equal to or greater than our observed difference in averages.

#### Host-associated morphological differences

To complement the genetic data, we asked whether there was evidence of morphological divergence among the three host lines. We focused specifically on ovipositor size and shape because ovipositor morphology has been linked to egg-laying success on thin-needled hosts (Bendall et al., 2017). If host-associated selection on needle width favors differences in ovipositor morphology and there is heritable variation in this trait, populations may exhibit host-associated phenotypic divergence. To control for the potential impact of rearing host on ovipositor morphology (i.e., plasticity), all females were reared on *P. banksiana* for at least two generations prior to dissection. We dissected, mounted, and imaged ovipositors from a total of 28 females (*n* = 10 Pitch females, 10 Shortleaf females, 8 Virginia females) as described in Bendall et al. (2017).

Briefly, a single lancet from each female was mounted on a glass microscope slide in an 80:20 permount:toluene solution. Each slide was imaged at 5x magnification and the ovipositor length and width were measured using the ZEN lite 2012 software package (Carl Zeiss Microscopy, LLC; Thornwood, NY). We compared ovipositors from each host using a geometric morphometric analysis, which computes shape differences while controlling for ovipositor size. We used IMAGEJ (v1.51; Schneider *et al*. 2012) to place a total of 30 landmarks defining the overall shape of each ovipositor (see Figure 3B) and transformed the position of each landmark into Cartesian coordinates. We aligned the landmarks of each ovipositor using a general Procrustes alignment in GEOMORPH (4.0.5; Adams and Otárola-Castillo 2013) implemented in R. Shape differences were visualized via a principle components analysis and assessed for significance using Procrustes ANOVA with sawfly line as a fixed factor. We also assessed differences in ovipositor length and width among sawfly lines using Type II ANOVAs via the *car* package.

### Assessment of Mechanisms Underlying IBE Pattern

#### Natural selection against immigrants

To determine whether sawflies derived from the three Arboretum hosts differ in their performance on different hosts, we reared the offspring of mated, lab-reared females from each host line on each of the three pine species. Briefly, after being mated to a same-host male, each female was placed in a mesh sleeve cage (25.4cm × 50.8cm) with a single seedling of one of the three host plant species (*P. echinata*, *P. virginiana*, or *P. rigida*; “no choice” scenario; see below). For each combination of host species and sawfly line (9 total), we performed 12-29 no-choice assays for a total of 144 no-choice assays.

No-choice cages were checked daily for eggs or until the female died, ultimately yielding between 5 and 10 families for each of the 9 sawfly-line x host-plant combinations. For each egg-bearing tree, the number of eggs laid was counted. Because *N. lecontei* females are born with their entire complement of eggs and tend to lay this entire complement in a single bout, the number of eggs laid on a tree would represent a female’s entire reproductive output. Egg-bearing trees were checked daily and watered as needed until larvae hatched (∼2 weeks). Because *N. lecontei* are gregarious and fare poorly when isolated, we reared siblings together in a single rearing box. Larvae were fed clippings from the same host species they hatched from *ab libitum* until cocooning. Cocoons were collected as they were spun, watered briefly to promote hardening, and weighed within 48 hours of collection.

If natural selection against immigrants contributes to IBE, we expected to see a significant difference among lines and/or a line-by-host interaction for one or more of the performance measures, with sawfly lines performing best on their host of origin (hereafter, source host). Therefore, we assessed performance differences between the lines in three ways: egg-to-cocoon survival, development time to cocoon, and cocoon weight.

To determine whether egg-to-cocoon survival rate on different hosts differed among the lines, we examined survival to cocoon at the colony level, where each colony was a family that consisted of a group of siblings with some number of eggs that hatched and survived to the cocoon stage and some number of eggs that did not. We excluded families for which no eggs hatched, as this is usually due to external variables (e.g., seedling death). We then used the *glmer* function (lmerTest package) to fit a mixed-effects logistic regression model with a logit link function to the survival data, with colony ID included as a random effect to account for differences among families in hatching success and larval performance unrelated to line of origin. Our model included line, rearing host, and a line-by-host interaction as fixed effects. To evaluate the significance of the two main effects and their interaction, we used a Type III ANOVA. For statistically significant main effects/interaction, we then used the *emmeans* package for *post hoc* pairwise comparisons. We used these comparisons to ask: (1) which pairs of lines differed in their overall egg-to-cocoon survival rates, (2) which pairs of hosts differed in their overall survival rates, and (3) for each line, whether there were differences in survival between pairs of hosts, as might be expected if each line performs best on its own source host. For each set of contrasts, we used a Benjamini-Hochberg correction for multiple comparisons.

For the subset of larvae that survived to the cocoon stage, we next asked whether the egg-to-cocoon development time on different hosts differed among the lines. Development time for each cocoon was calculated as the number of days between egg laying (the date on which the female was introduced into the sleeve cage) and cocoon spinning (the date the cocoon was collected in the rearing box, as recorded in our lab rearing logs). We used the *glmer* function to fit a mixed-effects Gamma regression model to the development time data, with colony ID included as a random effect and line, rearing host, and a line-by-host interaction as fixed effects. We evaluated the significance of the main effects and their interaction with a Type III ANOVA and used *emmeans* for *post hoc* comparisons as described above.

Finally, we asked if the sawflies that spun cocoons differed in weight. For females, which emerge with their full complement of eggs, cocoon weight correlates strongly with fecundity (Harper et al., 2016). For males, body size correlates with reproductive success (Glover et al., 2023). Because cocoon size is sexually dimorphic, we inferred the sex of each cocoon from weights and analyzed male and female cocoons separately. For each sex, we used the *lmer* command in lmerTest to fit a linear mixed model to the individual cocoon weights, with line and host as fixed effects and family as a random effect. We did not include a line-x-host interaction in the final models for male and female cocoon weights because the interaction term was not significant. After fitting the models, we evaluated significance of host and line effects with type II ANOVAs. We then used *emmeans* for *post hoc* pairwise comparisons among lines and hosts for each sex.

#### Sexual selection against immigrants

If sexual selection against immigrants contributes to host-associated differentiation, sawflies from each line should exhibit a decreased willingness to mate with individuals from other lines relative to their own line. To determine whether sawflies from each source host line are more likely to mate with individuals from the same host line, we conducted no-choice mating assays. We chose no-choice assays because one-on-one encounters most closely approximate mating in the wild (females often flee if approached by multiple males in the wild; Benjamin 1955). For each assay, a single virgin female was placed in a new, plastic 60mm x 12mm petri dish, and offered a virgin male from either the same line (Shortleaf x Shortleaf, Virginia x Virginia, and Pitch x Pitch) or a different line (Shortleaf x Pitch, Shortleaf x Virginia, and Virginia x Pitch, and the reciprocal crosses). To tease apart mating preferences from host preferences (which we evaluate below), we conducted mating assays in the absence of host material. To minimize the impact of inbreeding avoidance (Harper et al., 2016) on same-line mating assays, we obtained males and females from different propagation cages. Sets of six assays (3 same-line and 3 different-line) assays were recorded for 75 minutes with a Logitech or Microsoft web camera connected to a Lenovo Ideapad laptop. We switched the position of same-line and different-line pairings in each video to minimize positional biases. A total of 60 assays were performed for each type of pair, with 30 assays in each direction for different-line pairs. For example, to determine if there was sexual selection against immigrants between Shortleaf (SL) and Virginia (VA) lines, we set up 30 SL♀ x VA♂ crosses and 30 VA♀ x SL♂ crosses. In total, we recorded 360 no-choice mating assays. After filming, we reviewed the footage and recorded if mating occurred or not. We defined a mating event as an observed copulation lasting at least 60 seconds (Glover et al., 2023).

If there is sexual selection against immigrants, individuals should be less willing to mate with individuals from a different source host line. To test this prediction, we used the “glm” function in R with a “logit” link function to fit a binomial regression model to the mating outcome data, with female source (Shortleaf, Pitch, or Virginia), male source (Shortleaf, Pitch, or Virginia), and female source x male source interaction term as predictors. To evaluate the significance of the two main effects and their interaction, we used a Type III ANOVA.

#### Natural selection against hybrids

If natural selection acts against hybrids, offspring that are produced by crosses between different lines should have reduced fitness relative to offspring produced by parents from the same line. Ideally, hybrid performance would be compared to non-hybrid performance in all parental habitats, but for our study, the number of cross x host combinations was prohibitive (9 possible line combinations x 3 hosts = 27 treatments). Therefore, as a first step to evaluating the potential for reduced hybrid fitness to generate IBE in this system, we examined egg-to-cocoon survival of hybrids and non-hybrids on a single host, *Pinus banksiana*, not present at our study site. While this approach would not detect some potential sources of selection against hybrids, our rationale for using a non-native host was that we would still be able to detect reduced hybrid performance caused by genetic incompatibilities (e.g., due to physical linkage to or pleiotropic effects of divergently selected host use loci) or by maladaptive trait combinations (e.g., Bendall et al. 2017; Thompson et al. 2021) that cause hybrids to fare poorly on the non-native host.

To produce hybrid and non-hybrid larvae, we used mated females from our sexual isolation assays. After mating was observed, females were released into mesh cages with 8 *P. banksiana* seedlings. For each egg-laying female, we recorded the number of eggs laid, reared larvae on *P. banksiana* foliage as described above, and recorded the number of cocoons produced by each family. Due to variation in mating propensities and willingness to lay eggs, this resulted in an uneven number of families for the different crosses (P = Pitch, SL = Shortleaf, VA = Virginia): P♀ x P♂: *n* = 9, P ♀x SL♂: *n* = 10; P ♀x VA♂: *n* =10; SL♀ x P♂: *n* = 4; SL♀ x SL♂: *n* = 10; SL ♀x VA♂: *n* = 1; VA♀ x P♂: *n* = 11; VA♀ x SL♂: *n* =12; VA♀ x P♂: *n* = 9).

To test the prediction that hybrid families had reduced survival compared to non-hybrid families, we used the *glmer* function to fit a mixed-effects logistic regression model with a logit link function to the survival data, with maternal line, paternal line, and their interaction as fixed effects and colony ID as a random effect to account for differences among families in hatching success and larval performance unrelated to line of origin. To evaluate the significance of the two main effects and their interaction, we used a Type III ANOVA.

#### Dispersal bias via habitat preferences

Because *N. lecontei* mate on their host plant, divergence in host preferences may reduce gene flow between diverging host races, giving rise to a pattern of host-associated IBE. We therefore evaluated host preferences for all three lines in both no-choice and choice assays. No-choice assays were set up as described above (see “*Natural selection against immigrants*”). To determine whether lines differed in their willingness to lay eggs, we used the *glm* function to fit a logistic regression model to the binary (laid or did not lay) no-choice assay outcome data as a function of sawfly line, host plant, and their interaction. We used a Type III ANOVA to evaluate the significance of model terms and *emmeans* to perform *post hoc* tests.

For choice assays, individual females were released singly into 33cm x 33cm x 61cm mesh cages with two seedlings of each of two host species: the source host for the focal female and one of the two alternative host species. For choice assays, we used unmated females, which can lay unfertilized eggs that develop into haploid males and exhibit the same host preferences as mated females (Bendall et al., 2017). Due to constraints on space and availability of adult females and pine seedlings, we were unable to perform choice assays between non-source host pairs. In total, we conducted 193 choice assays, with roughly equal numbers across 6 experiments: Virginia females, *n* = 61 (*n* = 30 for *P. virginiana* vs. *P. echinata* assays; *n* = 31 for *P. virginiana* vs. *P. rigida* assays); Shortleaf females, *n* = 62 (*n* =31 for *P. echinata* vs. *P. rigida* assays; *n* = 31 for *P. echinata* vs. *P. virginiana* assays); Pitch females, *n* = 70 (*n* = 33 for *P. rigida* vs. *P. echinata* assays; *n* = 37 for *P. rigida* vs. *P. virginiana* assays). With these data, we asked whether females that laid eggs exhibited a preference for their source pine over alternative hosts in pairwise choice assays. Because each line had a unique set of choice assays, we analyzed each line separately. To determine whether a female from a particular line chose her source pine more often than expected by chance (50% based on equal numbers of source and non-source hosts offered), we used one-tailed binomial exact tests (*binom.test* function in R).

#### Habitat-related differences in phenology

To assess differences in the timing of the reproductive period between host lines, we tracked adult eclosion dates of all colonies collected in the field and returned to the lab in 2013 and 2014. Although *N. lecontei* typically has 2-3 generations per year in Kentucky, sawfly abundance varies across generations. Therefore, we focused our analyses on colonies that yielded at least 20 adults for the generation which we had sampling data for all three hosts available. In total, we collected eclosion data from 7 and 6 Virginia colonies, 5 and 3 Shortleaf colonies, and 3 and 3 Pitch colonies in 2013 and 2014, respectively. In each year, we tracked eclosion from the date of the first adult emergence. To quantify differences in adult phenology, we calculated pairwise estimates of temporal isolation (I) between populations following Feder et al. (1993):

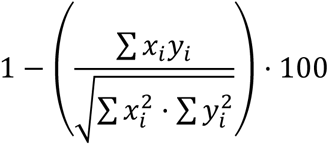

where *x_i_* and *y_i_* represent the proportion of the total number of live adults from host *x* or *y* on day *i*. We assumed an average lifespan of 5 days for females and 4 days for males based on field estimates (Benjamin 1955) and our own rearing experience. To assess patterns of eclosion between source hosts, we also compared the shape of cumulative eclosion curves for all adults collected from each of the three host species using bootstrapped Kolmogorov-Smirnov (KS) tests with 10,000 bootstraps with the *ks.boot* function from the *R* module MATCHING (v4.10-8; Sekhon 2008).

## Results

### Host plants are morphologically and chemically distinct

The three host plants differed from each other in all characteristics measured. For needle width, we found significant host-species and host-age effects, as well as a significant species-by-age interaction (Figure 2A; Supplemental Figure S2; Supplemental Table S6). For mature trees, all host species differed significantly from each other in needle width, with *P. echinata* having the thinnest needles and *P. rigida* the thickest. Except for *P. echinata* (*p* = 0.73), all seedlings were significantly thinner than their mature counterparts (*p* < 0.05).

**Figure 2.**
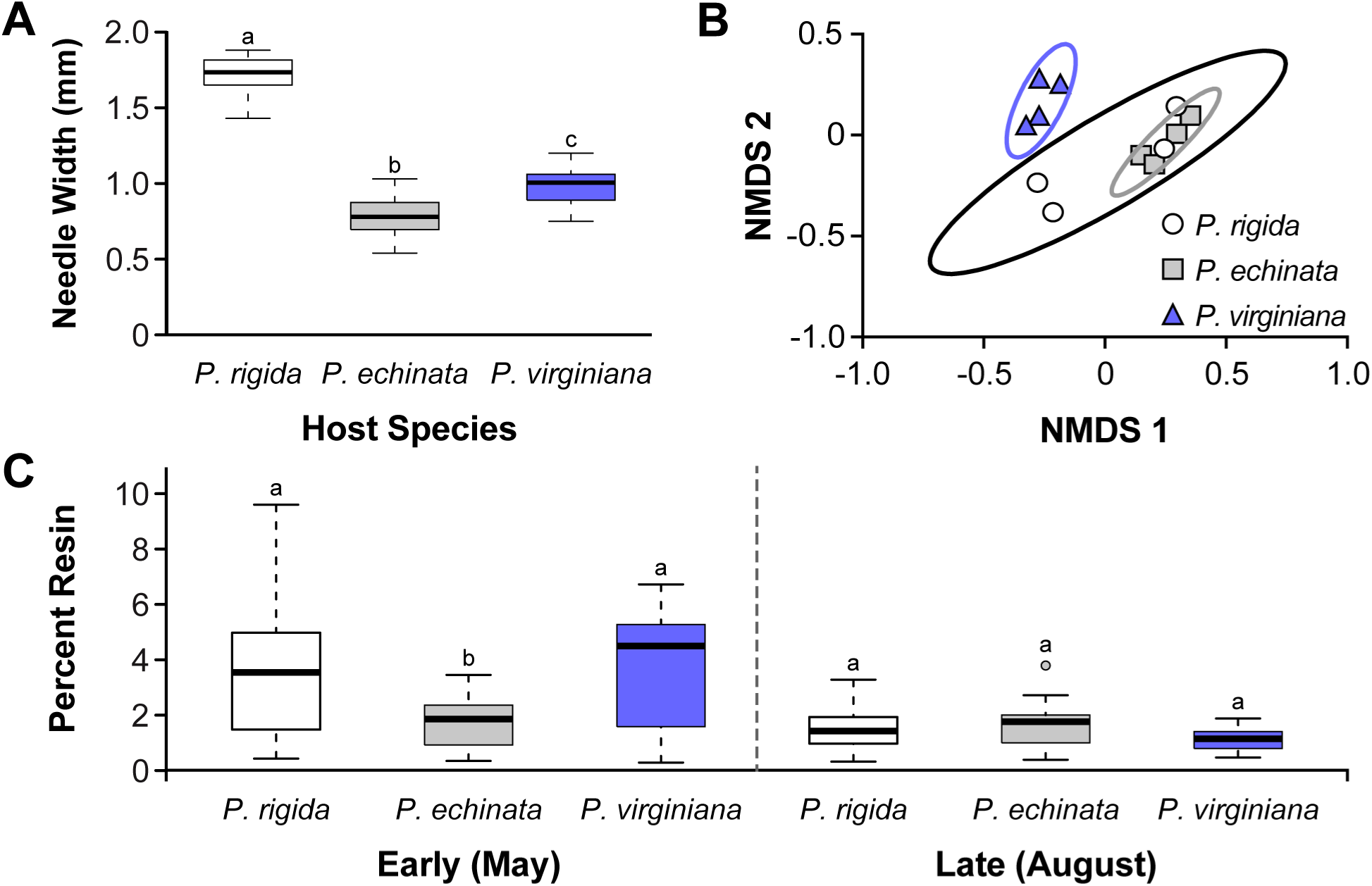
Host plant morphology, volatile profile, and resin content. A. Needle widths for mature trees sampled at the “Trail of Pines”. B. NMDS plot representing total volatile blend for the three host species. The volatile blend of *P. virginiana* is distinct, but those of *P. rigida* and *P. echinata* overlap. C. Resin content variation between the three hosts early (May) and late (August) in the sawfly season. For panels A and C, boxes represent interquartile ranges (median ± 2 SD), with outliers indicated as points; different letters represent comparisons that significantly differed in post-hoc comparisons.

**Figure 3.**
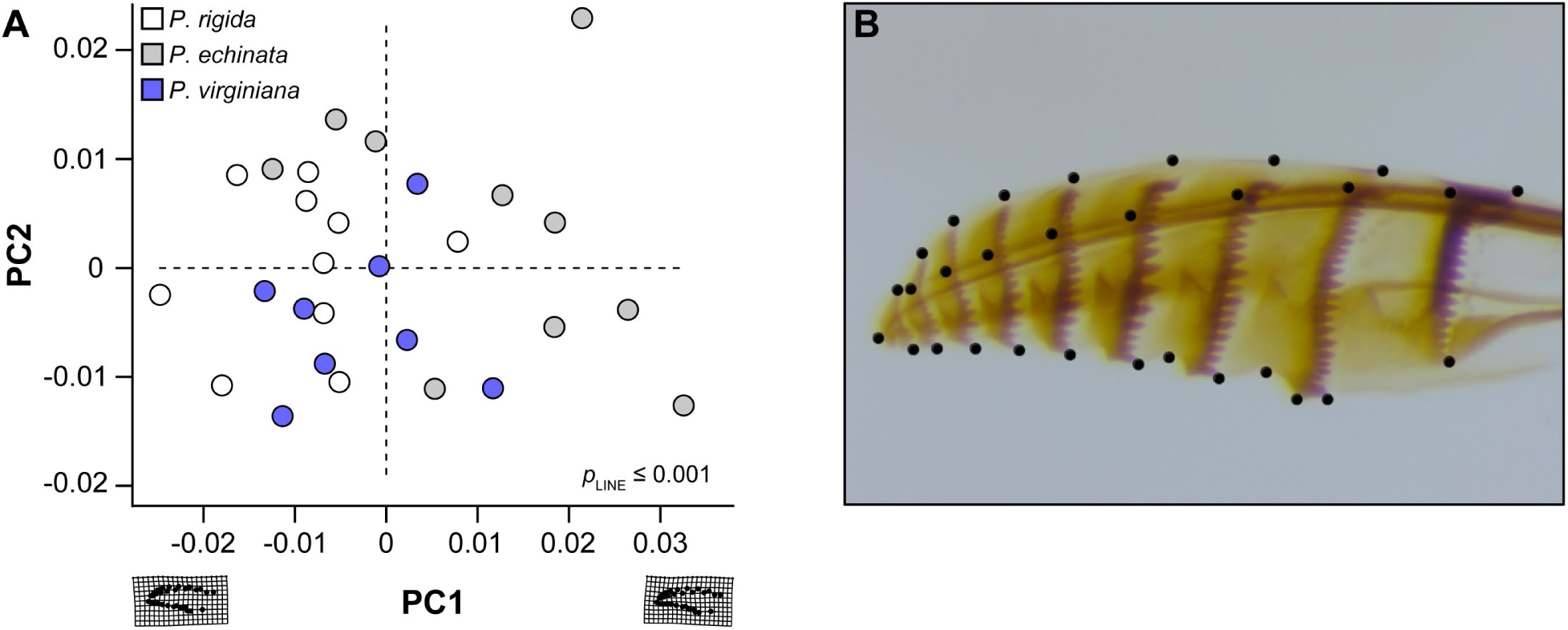
Variation in ovipositor morphology. A. Principle Components Analysis of overall ovipositor morphology, with Pitch females shown in white, Shortleaf females shown in grey, and Virginia females shown in blue. The warp grids demonstrate the change in shape along PC1. Shortleaf females have significantly differently shaped ovipositors than Pitch and Virginia females. B. Image of *Neodiprion lecontei* ovipositor showcasing the landmarks used in the analysis.

*P. virginiana* differed from *P. rigida* and *P. echinata* in total volatile profiles (Figure 2B). Variation in the volatile ratios appeared to drive the discrimination we observed in the total profile, as all abundant compounds (β-pinene, β-phellandrene, α-pinene, α-phellandrene, camphene, β-caryophyllene) and most minor compounds were produced by all three pines. Ratios of the abundant volatiles with linalool and β-caryophyllene varied by pine species (Supplemental Figure S3). *P. rigida* and *P. echinata* emitted different ratios of α-pinene, β-pinene, β-phellandrene, and β-caryophyllene against linalool (Supplemental Figure S3), which is noteworthy because these two species did not resolve based on total volatile profiles alone.

For resin content, there were significant host-species and sampling-month effects, and a significant interaction between host species and month (Figure 2C, Supplemental Table S7). Notably, *P. echinata* had less resin early in the sawfly season (May) than ether *P. rigida* or *P. virginiana*. Resin content declines in those hosts later in the season (August), such that no significant differences are observed. *P. rigida* and *P. virginiana* do not differ in resin content in May or August.

### Genomic data support IBE, but not historical isolation, among sawflies from different pines

Our ddRAD sequencing yielded 1.89 ± 2.35 (SD; standard deviation) million single-ended reads per individual; of which 1.88 ± 2.34 million survived quality filtering. After alignment, paralog filtering, and removal of putative PCR duplicates, an average of 0.95 ± 0.91 million alignments survived, and were formed into an average of 15,789 ± 7,271 RAD loci per individual with an average coverage of 45.67 ± 25.15x. These loci contained 33,674 SNPs. After removing seven individuals with high levels of missing data, two putatively haploid individuals, and enforcing a <30% missing data filter, the number of SNPs was reduced to 17,165. After applying the Hardy-Weinberg and minor allele frequency (MAF = 0.05) filters and subsampling to a single SNP per locus, our final dataset consisted of 6,759 SNPs.

Using this dataset, our evaluation of population structure selected *K* = 1 as the optimal number of clusters across all 100 independent runs, with CV error steadily increasing with *K* (Supplemental Figure S4). Values of *K* > 1 produced clustering solutions that were both unstable (multiple clustering solutions) and biologically uninterpretable (no clear assignment patterns). Furthermore, investigation of the clustering solutions offered under *K* = 2 and *K* = 3 revealed no meaningful structure between host lines (Supplemental Figures S5, S6). Overall, genome-wide pairwise F_ST_ between lines was modest, regardless of whether we considered all SNPs, genic SNPs only, or intergenic SNPs only (Table 1). Differentiation was significant between the Shortleaf and Virginia lines for all three SNP datasets. Although differentiation between Shortleaf and Pitch was similar to that observed for Virginia and Pitch, permutation tests were not quite significant. Finally, for both comparisons involving Virginia sawflies, genic differentiation was significantly higher than intergenic differentiation (Table 1).

**Table 1.** Pairwise F_ST_ for larvae collected on three different *Pinus* species. Pairwise F_ST_ values were computed using Hudson’s estimator and all SNPs, genic SNPs only, and intergenic SNPs only. P-values were obtained via permutation. Bolded values indicate significant differentiation (or significantly elevated differentiation at genic SNPs) at α=0.05.

| Host comparison | F<sub>ST</sub>, ALL SNPs<br>(P-value) | F<sub>ST</sub>, Genic SNPs<br>(P-value) | F<sub>ST</sub>, Intergenic<br>SNPs<br>(P-value) | P-value, F<sub>ST</sub><br>genic vs. F<sub>ST</sub><br>intergenic |
| --- | --- | --- | --- | --- |
| Pitch x Shortleaf | 0.0132<br>(P = 0.0571) | 0.0117<br>(P = 0.0800) | 0.0144<br>(P = 0.0551) | P = 0.9316 |
| Pitch x Virginia | 0.0047<br>(P = 0.2706) | 0.0062<br>(P = 0.1906) | 0.0030<br>(P = 0.4146) | **P < 0.0001** |
| Shortleaf x<br>Virginia | **0.0144**<br>**(P = 0.0168)** | **0.0151**<br>**(P = 0.0109)** | **0.0131**<br>**(P = 0.0340)** | **P = 0.0110** |

### Sawfly lines from different pines differ in ovipositor shape

Ovipositor shape differed among sawfly lines (Figure 3A; Supplemental Table S8). Shortleaf females differed from both Virginia and Pitch females; but Virginia and Pitch females did not differ in ovipositor shape. We did not detect differences in ovipositor length or width among sawfly lines (Supplemental Table S9).

### Evidence of natural selection against immigrants

For survival to cocoon, there was a significant effect of sawfly line and a significant line-by-rearing host interaction (Figure 4A, Supplemental Table S10). Overall, the Shortleaf line had higher egg-to-cocoon survival than the other two lines. Within the Pitch and Shortleaf lines, we did not detect any survival differences based on rearing host. However, the Virginia line had significantly reduced survival when reared on *P. rigida* compared to *P. virginiana* or *P. echinata*. For development time, all model terms (line, host, host-by-line interaction) were significant (Figure 4B; Supplemental Table S11). Notably, each sawfly line tended to develop fastest when reared on its original source host (i.e., Virginia sawflies developed fastest when reared on *P. virginiana*). Averaged across hosts, the Shortleaf line tended to develop faster than the other two lines. Averaged across lines, sawflies tended to develop slower on *P. rigida* than on the other two hosts. For female cocoon weight, we found significant sawfly-line and rearing-host effects, but their interaction was not significant (Figure 4C, Supplemental Table S12). Overall, Shortleaf females weighed less than Pitch and Virginia females. Additionally, regardless of sawfly line, females reared on *P. echinata* tended to weigh less than females reared on other pines. Male cocoon weight results were qualitatively very similar to those for female cocoon weight (Supplemental Figure S7, Supplemental Table S13): there were significant line and host effects, and Shortleaf males weighed less than males from other lines. One difference, however, was that males reared on *P. virginiana* were significantly heavier than those reared on other hosts. Together, these data indicate that there are substantial differences in larval performance traits within and among sawfly lines and host plants. These data also indicate that possible costs to immigrants (i.e., sawflies that choose a host species that differs from their source host) include reduced survival of eggs to cocoon (Figure 4A; Virginia line) and longer development times (Figure 4B; all lines).

**Figure 4.**
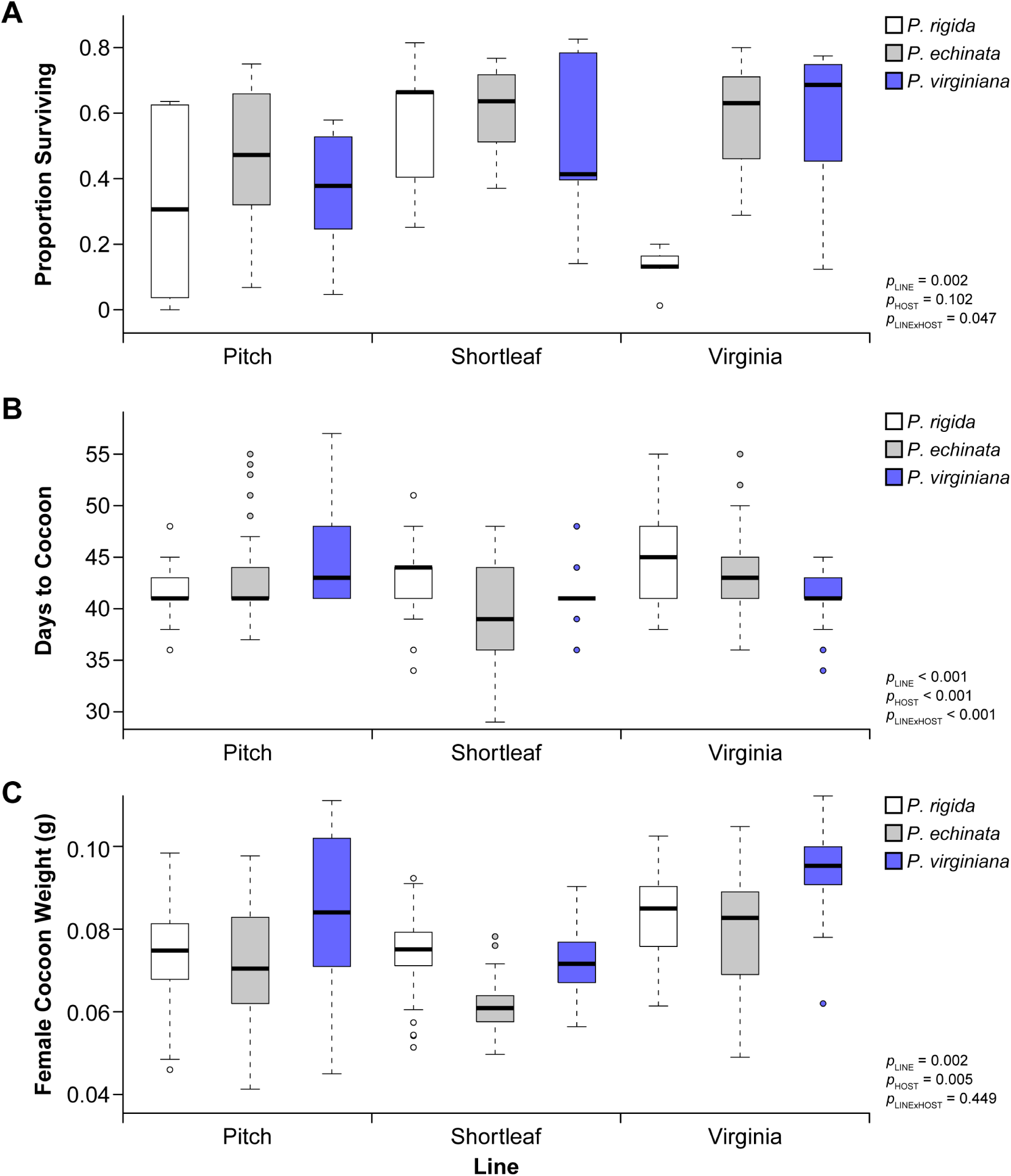
Evidence of selection against immigrants. A. Proportion of larvae surviving from egg to cocoon stage. B. Development time from egg to cocoon. Sawflies tended to develop fastest on their source host. C. Female cocoon weight. Shortleaf females had the lowest cocoon weights, and females reared on *P. echinata* tended to be smaller than those reared on other hosts. For all panels, boxes represent interquartile ranges (median ± 2 SD), with outliers indicated as points. Model statistics are given for each panel, with statistics for post-hoc pairwise comparisons given in Supplemental Tables 10-12.

### No evidence of sexual selection against immigrants or natural selection against hybrids

Neither female line nor male line had any effect on mating outcomes in no-choice assays (Supplemental Table S14). Likewise, the combination of male and female lines (male x female interaction) did not affect mating outcome. Together, these results indicate that females and males from all lines were equally willing (or unwilling) to mate, regardless of whether they were paired with an individual that came from the same or a different host species. As all individuals used in this assay were reared on *P. banksiana*, however, we cannot rule out the existence of mate discrimination using a host-odor related cue in natural populations.

When we reared offspring of mated pairs on a non-native host, we found no effect of maternal line, paternal line, or their interaction on offspring survival to cocooning (Supplemental Table S15). The hybrid offspring produced by the different-host pairings did not have any obvious reduction in survival compared to non-hybrid offspring, although we note that sample sizes for some cross types were small. In addition, there could be other sources of reduced hybrid fitness (e.g., reduced survival on other host species, reduced adult reproductive success) our approach could not detect.

### Some evidence of biased dispersal via habitat preferences

In no-choice assays, the host plant offered did not influence a female’s likelihood of ovipositing, nor was there a significant line-by-host interaction (Supplemental Table S16). However, there was a significant effect of the female’s line on overall willingness to lay eggs (Figure 5A). Specifically, Shortleaf females were more likely to oviposit than Pitch and Virginia females. In choice assays, only Virginia females were significantly more likely to lay their eggs in their host (*P. virginiana*) than a non-source host (*P. echinata* or *P. rigida*) (Figure 5B; Supplemental Table S17). Although we could not compare choice assays directly because they were set up differently for different lines, we note that the proportions of females from each line that laid eggs in choice assays were very similar to those observed in no-choice assays: 81% of Shortleaf females, 54% of Pitch females, and 61% of Virginia females laid eggs in choice assays. Overall, our host preference assays reveal that Pitch and Virginia females are more reluctant to lay on pine seedlings than Shortleaf females and that Virginia females prefer their source host.

**Figure 5.**
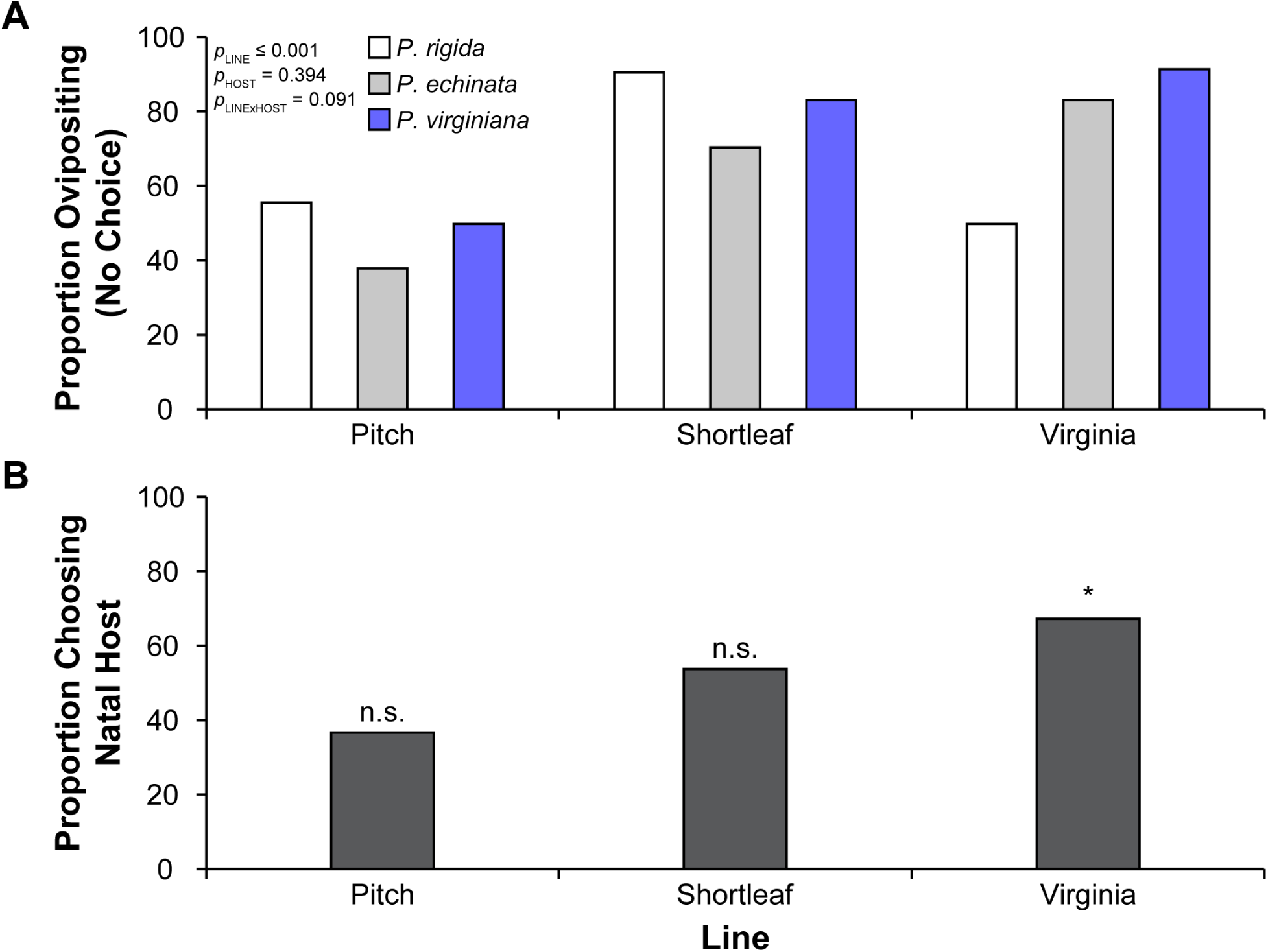
Evidence of dispersal bias amongst sawfly lines. A. Proportion of females ovipositing in no-choice assays. P-values for model terms are given; post-hoc comparisons among sawfly lines are in Table S16. B. Proportion of females choosing their source host over non-source hosts in choice assays. Only Virginia females demonstrated a preference for their source host (*P. Virginiana*). For panel B, n.s. indicates p > 0.05; * indicates p < 0.05.

### Evidence of host-related phenology differences

Although patterns of eclosion varied between host plants and between years, there was at least partial temporal isolation between the lines in each year (Figure 6; Table 2). In 2013, all sawfly lines differed significantly in their eclosion pattern, although the Pitch and Virginia lines were less strongly isolated than the other comparisons. In 2014, the Shortleaf and Virginia lines did not significantly differ in eclosion pattern and were also less isolated than in 2013. Conversely, although their eclosion patterns significantly differed in both years, the Pitch and Virginia lines were more strongly isolated in 2014 than in 2013. The Pitch and Shortleaf lines significantly differed in eclosion patterns and were strongly isolated in both years.

**Figure 6.**
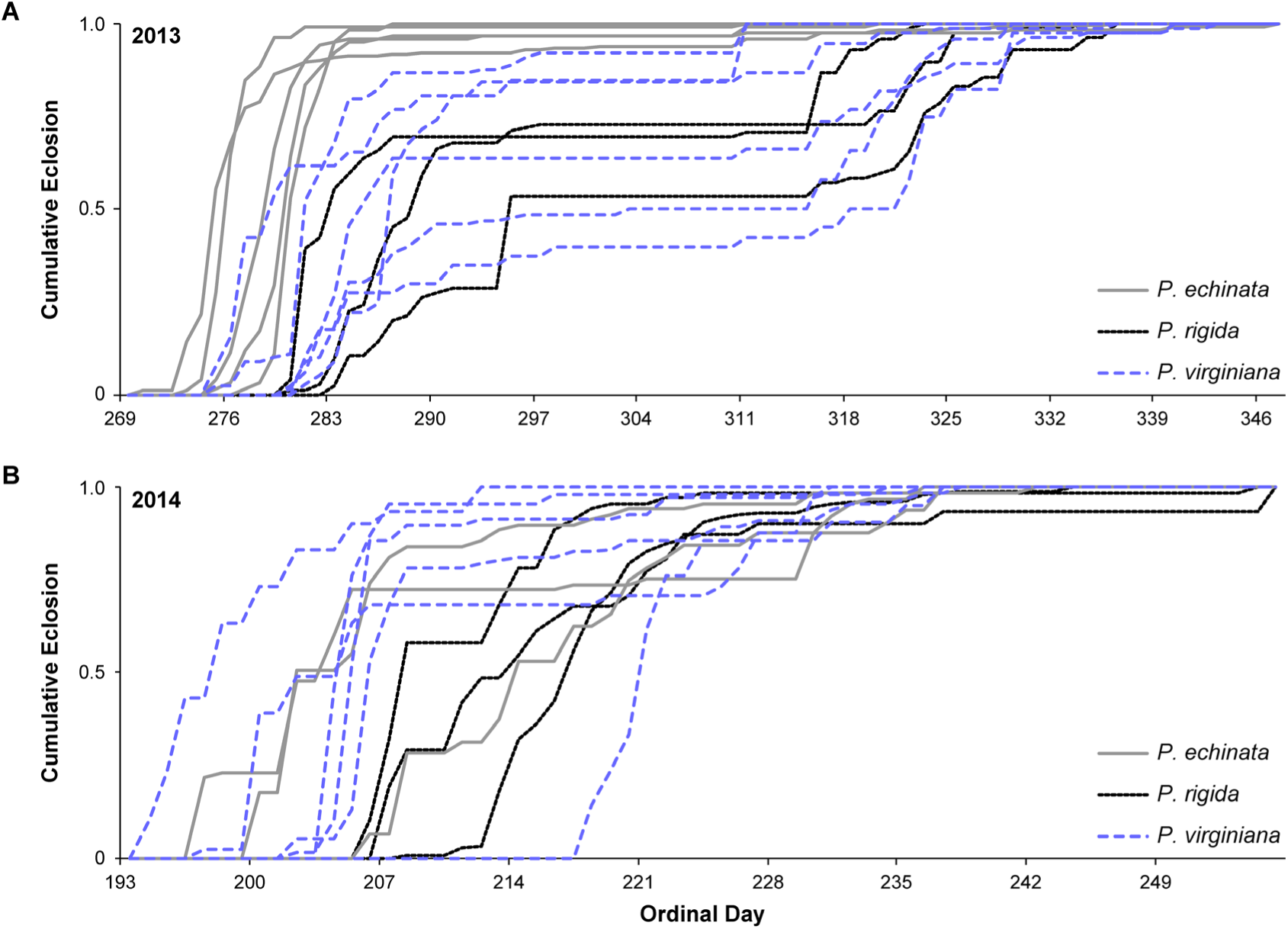
Evidence of host-related phenological differences. Cumulative eclosion curves for individual families with at least 20 eclosing adults from (A) 2013 and (B) 2014. Colonies from *P. echinata* are shown in solid grey lines, from *P. rigida* in solid black lines, and *P. virginiana* in dashed blue lines. In 2013, all hosts differed significantly in their eclosion patterns. In 2014, Shortleaf and Virginia lines were no longer isolated (see Table 2).

**Table 2.** Phenological differences in adult eclosion patterns. For each pairwise comparison, the pairwise I values (upper number) and p-values from KS tests (lower number) are given. Both evaluate differences in the cumulative eclosion curve per source host.

|  | Pairwise I |  |
| --- | --- | --- |
|  | 2013 | 2014 |
| Pitch x Shortleaf | **0.8155**<br>(p < 0.0001) | **0.7026**<br>(p = 0.01891) |
| Pitch x Virginia | **0.4851**<br>(p = 0.0119) | **0.7188**<br>(p = 0.01891) |
| Shortleaf x Virginia | **0.7004**<br>(p < 0.0001) | 0.3683<br>(p = 0.5308) |

## Discussion

In many species, including the redheaded pine sawfly *Neodiprion lecontei*, genetic differentiation between populations increases as their environments become more dissimilar (Bagley et al., 2017; Sexton et al., 2014; Shafer & Wolf, 2013; Wang et al., 2013). To better understand how a pattern of isolation by environment evolves, we characterized patterns and mechanisms of divergence in a sympatric population of *Neodiprion lecontei* that recently colonized three pine hosts. We first characterized differences in needle structure, volatile profiles, and defensive chemistry among the three pine host plants that could generate divergent selection pressures, bias dispersal patterns, and promote phenological differences. Next, we evaluated patterns of genetic differentiation among sawflies collected from the different hosts, finding modest differentiation between the three lines consistent with recent colonization and *in situ* divergence. Our laboratory assays support three potential mechanisms generating IBE in *Neodiprion lecontei*: natural selection against immigrants, biased dispersal, and host-related differences in phenology. However, we find no evidence of sexual selection against immigrants or selection against hybrids in this system. Overall, our results suggest that different mechanisms can contribute to IBE between populations even when they belong to the same species and occupy the same geographic location. Below, we consider limitations of our data and discuss how these results impact our understanding of *Neodiprion* divergence and, more generally, how environmental differences shape patterns of genetic variation in nature.

### Rapid emergence of phenotypic differentiation and IBE

Our population structure analyses suggest that our focal population represents a single genetic cluster, rather than multiple, already-diverged lineages. As a result, the host-associated phenotypic and genetic differentiation we observed most likely arose *in situ* and rapidly (within <40 generations). Rapid phenotypic adaptation to host plants has also been documented in other insect systems (e.g., Singer et al. 1993; Thompson 1998; Sousa et al. 2019), often following introduction of pest insects or of novel host plants. For example, within the past 50-80 years, several host races of the American soapberry bug (*Jadera haematoloma*) have evolved, with the mouthpart (“beak”) length corresponding to the size of the host’s fruit (Carroll & Boyd, 1992; Comerford et al., 2022). These host races also have faster development time and greater survival on their novel hosts (Carroll et al., 1997, 1998).

In our study population, patterns of host-associated phenotypic differentiation among the three pine hosts involved multiple phenotypic traits, including differences in host preference (Figure 5), larval development on different host plants (Figure 4B), body size (Figure 4C), and ovipositor morphology (Figure 3A). Although phenotypic differences can result from plastic responses to rearing host (e.g., Görür 2003; Pfennig et al. 2010), we measured all traits in individuals that had been reared in a common lab environment and on the same non-source host plant (*P. banksiana*), suggesting the observed phenotypic differences were heritable.

Perhaps the most striking morphological differences were between sawflies that originated from *P. echinata* (shortleaf pine) and sawflies that originated from the other two pine species. Shortleaf females had differently shaped ovipositors (Figure 3A) and Shortleaf cocoons—which predict adult body size—were smaller than cocoons from Pitch and Virginia lines (Figure 4C). Notably, although all three host species differ in needle width, *P. echinata* has the thinnest needles (Figure 2A). Our finding that *N. lecontei* adults from the thinnest-needled host were smaller than adults from other hosts is consistent with recent work demonstrating concordant host needle-width and adult body-size clines in *N. lecontei* (Glover et al., 2023). Also, our finding that these morphological differences are maintained in sympatry is consistent with experimental work demonstrating that thin-needled pines impose strong selection favoring smaller ovipositors (Bendall et al., 2017) and smaller eggs (Glover et al., 2023) in *Neodiprion* females.

In addition to the observed phenotypic differences, we also observed modest genetic differentiation (F_ST_)—or isolation by environment (host)—between Shortleaf lines and the other two lines. However, differentiation between Shortleaf and Pitch lines was not quite significant (Table 1), possibly due to a smaller sample size for this comparison. Given the observed phenotypic differentiation among the three sawfly lines, the observed genetic differentiation could stem in part from divergent selection on loci encoding traits that impact performance on different pines. In support of this hypothesis, differentiation at genic loci (putatively selected) was significantly higher than differentiation at intergenic loci (putatively neutral) for two of three pairwise comparisons (Table 1). However, we also observed host-related differentiation in intergenic regions, suggesting that neutral differentiation was facilitated by reduced effective migration between Shortleaf sawflies and sawflies from other hosts (Table 1).

Although little is known of the speed at which IBE can emerge, evidence from multiple invasive species indicates that IBD can be generated quickly. For example, isolation by distance was detected within 15 years of invasion amongst Asian tiger mosquitoes collected in villages in the Torres Straight Islands in Australia (Schmidt et al., 2021). The IBD pattern was also detected in western corn rootworm during both its initial invasion of and after establishment and spread through Southern Europe (Lemic et al., 2015). In the pine-sawfly family Diprionidae, IBD has been detected in *Diprion similis*, a species that invaded eastern North America in the early 1900s (Davis et al. In Press). Here, we report the rapid emergence of IBE at a single site, likely within no more than 40 generations. Given the potentially important role that selection plays in generating IBE, it is possible that IBE tends to emerge more quickly than IBD. However, much more work is needed to evaluate the relative speed with which these different patterns of population structure tend to emerge.

### Multiple mechanisms contribute to IBE in pine sawflies

Our work supports at least three mechanisms contributing to IBE in pine sawflies. First, our larval performance experiments suggest there may be natural selection against immigrants on different pines. For example, all lines developed most quickly on their source host (Figure 4B). Under field conditions, prolonged development on alternative hosts could impose selection against immigrants via increased exposure to a large community of pathogens, parasitoids, and predators (see Holling 1959; Hanski and Parviainen 1985; Olofsson 1987; Wilson et al. 1992; Forbes et al. 2018). Natural selection against immigrants was also evident for the Virginia line, which had reduced survival to the cocoon stage when reared on a non-source host, *P. rigida* (Figure 4A). Finally, although we found no evidence to indicate that sawflies produced larger cocoons when reared on their source host, Shortleaf males and females tended to have smaller cocoons than males and females from the other lines (Figure 4C; Supplemental Figure S7). Although lower cocoon weights are associated with reduced fecundity in females (Harper et al., 2016), reduced body size and egg size is also associated with increased hatching success on thinner needled pines (Bendall et al., 2017; Glover et al., 2023). Thus, smaller body sizes could explain why the Shortleaf line had the highest egg-to-cocoon survival rates in our laboratory assays (Figure 4A), which used seedlings for oviposition hosts. Notably, seedlings have even thinner needles than any of the mature hosts (Figure S2).

However, because we did not directly evaluate hatching success in our survival assays, additional experiments are needed to determine the relative contribution of adult female oviposition traits and larval feeding traits to immigrant inviability. To more closely approximate selection in the field, such experiments would ideally use mature host plants for oviposition rather than seedlings. More generally, because laboratory assays can miss important sources of divergent selection and immigrant inviability (Hatfield & Schluter, 1999; Kimball et al., 2008; Rundle & Nosil, 2005), field-based diet manipulations are a high priority for future work.

Second, we also found evidence of a potential dispersal bias: in both choice and no-choice assays, Virginia females were more likely to oviposit on *P. virginiana* than on other hosts (Figure 5B). Interestingly, *P. virginiana* has the most distinct volatile profile of the three hosts at the Arboretum (Figure 2B), offering a potential explanation for why the Virginia line was the only sawfly line to demonstrate a strong host preference. Additionally, females from the Virginia line were especially reluctant to oviposit on *P. rigida* (Figure 5A), a host on which this line had reduced survival (Figure 4A). These observations suggest that divergent host preferences in the Virginia line evolved via natural selection. Consistent with this hypothesis, all pairwise comparisons involving the Virginia line had significantly higher F_ST_ at genic regions compared to intergenic regions (Table 1). Because *Neodiprion* sawflies mate on their host plant (Benjamin, 1955; Coppel & Benjamin, 1965; Knerer, 1984), divergent habitat preferences also have the potential to reduce gene exchange among hosts. Host fidelity—the tendency of individuals to reproduce on their natal host type—has long been thought to facilitate sympatric speciation (Feder et al., 1994; Hirai et al., 2006; Wood et al., 1999). Similarly, the reduced gene flow that results from divergent habitat preferences may frequently contribute to IBE.

Third, across three years, we found evidence of strong, but sometimes variable temporal isolation among the three hosts (Figure 6; Table 2). Overall, Pitch and Shortleaf lines were the most consistently and strongly isolated in terms of phenology, with Shortleaf adults tending to emerge early and Pitch adults tending to emerge late. This partial temporal isolation is likely to reduce gene exchange, enabling neutral regions of the genome to diverge via drift. Consistent with this prediction, the Pitch x Shortleaf comparison had the highest intergenic F_ST_ of the three pairwise comparisons, with no detectable difference between genic and intergenic regions (Table 1). Like host fidelity, temporal isolation is thought to play an important role in promoting speciation in the absence of geographic barriers (reviewed in Taylor and Friesen 2017). Not only is temporal isolation a particularly effective barrier to gene exchange (Abbot & Withgott, 2004; Feder et al., 1993, 1994), variation in abiotic and biotic selection pressures among habitats often generate divergent selection on reproductive timing (Burban et al., 2020; Hood et al., 2019; Santos et al., 2011; Svensson et al., 2005; Thomas et al., 2003). For these reasons, temporal isolation may be an especially common mechanism generating IBE.

Although temporal isolation appears to contribute to IBE in *N. lecontei*, we cannot determine from our data—which were collected from field-caught mid-late instar larvae reared to adulthood in the lab—whether variation in adult emergence times is due to genetic variation, rearing environment, or both. One potential explanation for differences in adult emergence timing is that these differences evolved via natural selection to optimize timing for different hosts (e.g., Feder et al. 1993, 1994; Feder and Forbes 2010). Although we do not yet know which host cues *N. lecontei* females use when selecting host plants for oviposition (but see Tisdale and Wagner 1991; Björkman et al. 1997), we do know that pines as a whole (Nerg et al., 1994) and the specific trees at the Trail of Pines (Figure 2C) vary seasonally in resin content. Pines also vary seasonally in moisture levels (C. E. Van Wagner, 1967) and volatile profile (Geron & Arnts, 2010). If the seasonal variation in host quality differs between the three host species, selection could favor different peak emergence times among sawflies using different pines. Finer grained analysis of seasonal variation in host quality, impact on sawfly reproductive success, and heritability of adult emergence times are required to evaluate this hypothesis. Eclosion differences could also be generated via plasticity in development time. Developmental plasticity is well documented in insects (see Nylin and Gotthard 2003), with many examples demonstrating that larvae develop at different rates when reared on different diets. Indeed, our results indicate that rearing host affects developmental timing, with evidence of a genotype-by-host interaction as well (Figure 4B). Confirming that differences in the speed of egg to cocoon development give rise to differences in adult eclosion timing in the field will require additional field surveys.

Unlike immigrant inviability, biased dispersal, and temporal isolation, we did not find any evidence of sexual selection against immigrants: mating outcomes did not differ between same-host and different-host pairs in no-choice mating assays (Supplemental Table S14). By contrast, a recent experiment using the same mating assay design revealed sexual isolation between *N. lecontei* and sister-species *N. pinetum*, largely stemming from strong size-based assortative mating within and between species (Glover et al., 2023). Despite some size differences among the three lines in this experiment (Figure 4C; Supplemental Figure S7), these differences were apparently insufficient to produce strong assortative mating by host in our assays. One limitation of our mating assays, however, is that all individuals were reared on the same host plant, potentially minimizing size differences arising because of rearing host. Rearing host clearly influences body size in *N. lecontei* (Figure 4C; Supplemental Figure S7), and work in other insect systems demonstrates that dietary influences on body size can affect mating outcomes (Forister & Scholl, 2012). Rearing diet could also influence assortative mating via affecting adult pheromone composition, cuticular hydrocarbon profiles, or chemical-based mating preferences (e.g., Conner et al. 1990; Gosden and Chenoweth 2011; Darragh et al. 2019). Finally, in nature, mating typically takes place on the host plant, so it’s possible that the presence of host material interacts with other behavioral and chemosensory cues to influence mating outcomes (e.g., Sattman and Cocroft 2003; Liao et al. 2016). Thus, to rule out sexual selection against immigrants, additional experiments are needed.

We also did not find evidence of reduced hybrid viability in our assays, as might be expected if there were partial intrinsic postzygotic isolation between the three lines. We note, however, that we did not measure fertility, fecundity, or mating success of hybrids. Also, perhaps the biggest limitation of our hybrid assays was that we did not evaluate host-based sources of reduced hybrid fitness. As ecologically dependent selection against hybrids has been noted in many other insect systems [Servedio 2004; e.g., *Rhagoletis* flies (Linn et al., 2004), *Timema* walking sticks (Sandoval 1994a,b), and pea aphids (Via et al., 2000)], it may contribute to IBE in *N. lecontei*. For example, hybrid females between *N. lecontei* and its sister species *N. pinetum* have mismatched host preferences and egg-laying traits that drastically reduce oviposition success (Bendall et al., 2017). Although the magnitude of differences in host preference and ovipositor differences revealed in this study are modest compared to interspecific differences, mismatches in these traits may nevertheless offer at least one mechanism by which hybrid females at the site could have reduced fitness compared to non-hybrids.

## Conclusion

Like many taxa investigated to date (Gray et al., 2014; Prunier et al., 2017; Sexton et al., 2014; Shafer & Wolf, 2013; Weber et al., 2017), genetic differentiation among populations of *Neodiprion lecontei* correlates with both geographic distance (IBD) and host use (IBE). Here, we take advantage of a sympatric population of *N. lecontei* on three hosts to explore potential mechanisms producing IBE. Our analyses reveal both that a pattern of IBE can emerge very rapidly—in our case, in tens of generations—and that multiple mechanisms—including immigrant inviability, dispersal bias, and temporal isolation—are likely important to generating IBE. These two observations may be related: perhaps IBE emerges quickly precisely because there are multiple mechanisms simultaneously reducing effective migration rate between habitats. Still, much work remains to better understand IBE in this system. Additionally, comparative work across diverse taxa are needed to evaluate the relative importance of different IBE mechanisms and to test the hypothesis that IBE evolves more quickly than IBD. While such work is labor-intensive, it is essential for better understanding patterns of genetic variation in nature.

## Author contributions

RKB and CRL designed the research. RKB, MNH, JF, JW, JWT2, CJF, VCS, and CRL performed the research. RKB, CJF, VCS, and CRL analyzed the data. RKB and CRL wrote the manuscript with input from all authors.

## Acknowledgements

We thank the staff of the University of Kentucky’s Arboretum and State Botanical Gardens, especially Todd Rounsaville, for allowing us to collect sawflies, clip branches, and conduct research at the site. We also thank the members of the Linnen lab for their assistance with monitoring the site and rearing of pine sawflies used in these experiments. We thank Dylan P’Simer for assistance with the resin extraction protocol. We also thank Vitor Sousa for helpful discussions of F_ST_ analyses, as well as for providing several R scripts. This work was supported by the National Science Foundation (DEB-1257739 and DEB-CAREER-1750946 to CRL, IOS-2101059 and IOS-1656625 to CJF) and USDA-NIFA (predoctoral-fellowship 2015-67011-22803 to RKB), the BIO5 Institute (CJF), and the University of Kentucky’s Ribble Travel Grant (to JWT2). For computing resources, we thank the University of Kentucky Center for Computational Sciences and the Lipscomb High Performance Computing Cluster.

## Data accessibility statement

All datasets, input files, R codes, and R scripts used in genetic and IBE assays described in the study will be made available in standard formats on DRYAD (doi TBD). Short read data will be available on the NCBI Short Read Archive.

## Supplemental Information

- Figures

- Figure S1. Appearance and organization of the study site.
- Figure S2. Comparison of needle width in mature trees and seedlings.
- Figure S3. Ratios of abundant volatiles across three pine species.
- Figure S4. C.V. error plots for ADMIXTURE runs.
- Figure S5. Clustering solutions for K = 2.
- Figure S6. Clustering solutions for K = 3.
- Figure S7. Comparison of male cocoon weights.
- Tables

- Table S1. Collection information for resin content measurement.
- Table S2. Usage for each *Neodiprion lecontei* colony.
- Table S3. Sequencing information for each specimen.
- Table S4. Sequences for ddRAD adapters.
- Table S5. PCR primers for amplification of ddRAD libraries.
- Table S6. Statistical results for needle width comparisons.
- Table S7. Statistical results for resin content comparisons.
- Table S8. Statistical results for ovipositor morphology analysis.
- Table S9. Statistical results for ovipositor length and width comparisons.
- Table 10. Statistical results for larval survival comparisons.
- Table S11. Statistical results for development time comparisons.
- Table S12. Statistical results for female cocoon weight comparisons.
- Table S13. Statistical results for male cocoon weight comparisons.
- Table S14. Statistical results for mating outcome comparisons.
- Table S15. Statistical results for hybrid survival outcome comparisons.
- Table S16. Statistical results for no-choice host preference assays.
- Table S17. Statistical results for choice host preference assays.

