## Supplemental Figures for "Multiple mechanisms contribute to isolation-by-environment in the redheaded pine sawfly, *Neodiprion lecontei*"

**Figure S1. Appearance and organization of the study site.** The “Trail of Pines” is a ~130m transect of pine trees located in The Arboretum State Botanical Garden of Kentucky. It contains several large pine trees of three species native to Kentucky which were planted in the 1990’s. Panel A shows the general placement of the mature pines at the site, with pitch pines (*Pinus rigida*) shown in white, shortleaf pines (*P. echinata*) shown in grey, and Virginia pines (*P. virginiana*) shown in blue. The trees at the site are in very close proximity to each other, with the branches of adjacent trees often intermingling, as shown in panel B.

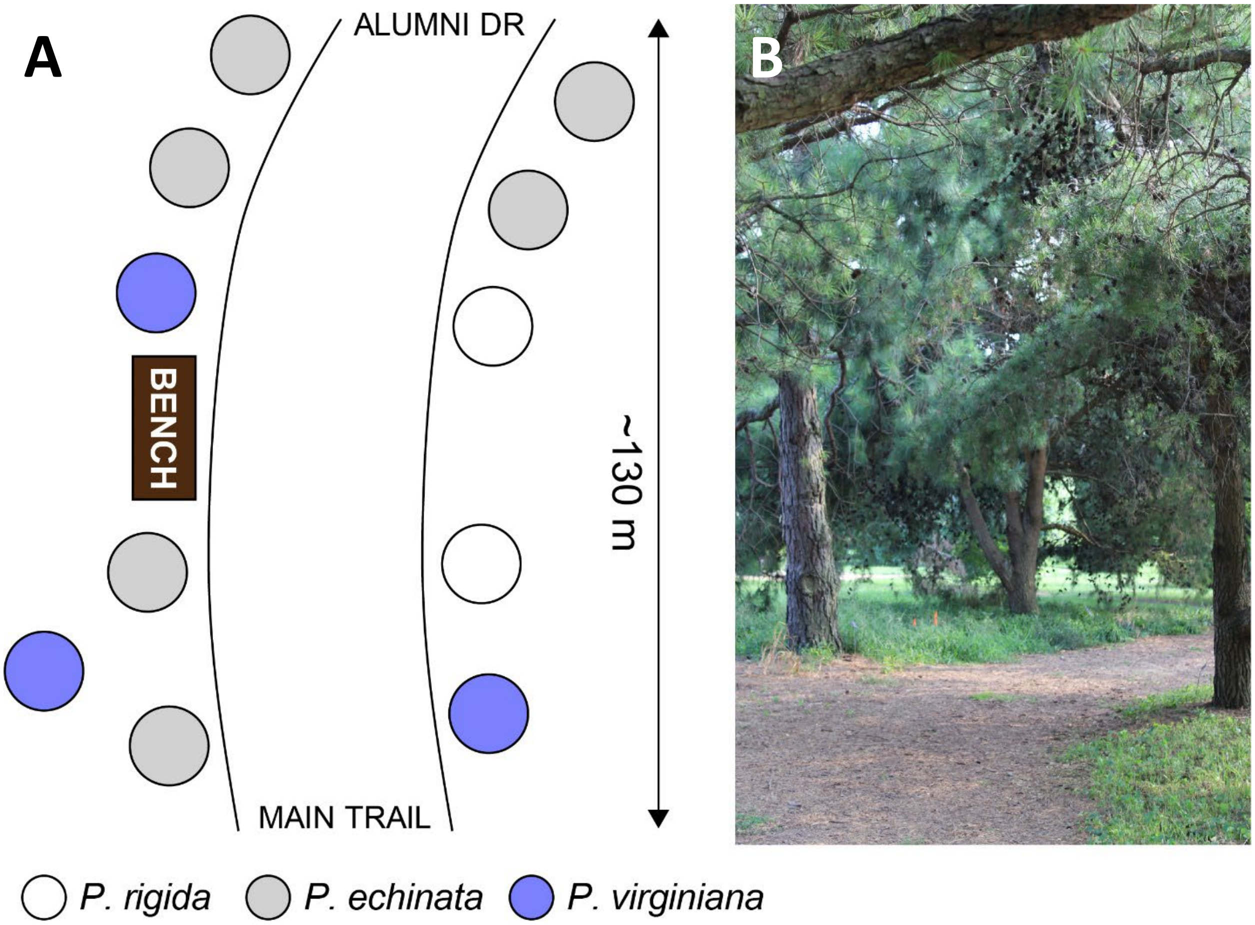

**Figure S2. Comparison of needle width in mature trees and seedlings.** Lower case letters represent significant differences in pairwise post-hoc comparisons between mature trees. Upper case letters represent significant differences in pairwise post-hoc comparisons between seedlings. The bars represent comparisons between needles of mature trees and seedlings for each host (*ns* – p > 0.05; * – p < 0.05; *** – p < 0.0001). All mature trees are significantly different in needle width. In seedlings, only *P. rigida* and *P. echinata* had significantly different needle widths. Seedlings of *P. rigida* and *P. virginiana* had significantly thinner needles than their mature counterparts, but no such difference existed within *P. echinata*.

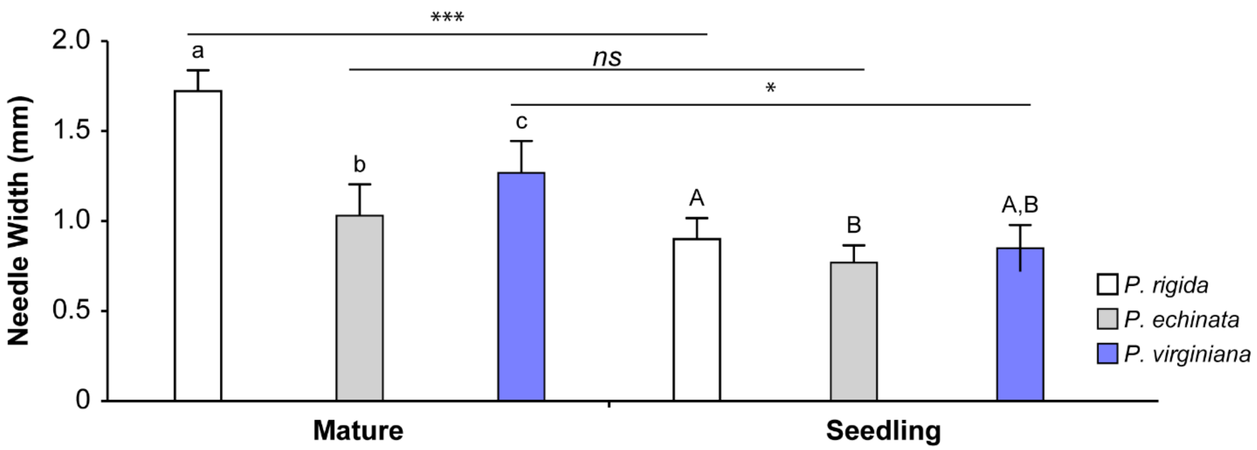

**Figure S3. Ratios of abundant volatiles across three pine species.** Box plots of select pairwise ratios between α-pinene, β-pinene, linalool, α-phellandrene, β-phellandrene, β-caryophyllene, and camphene. Dots, triangles, and squares are individual data points for *P. rigida*, *P. echinata*, and *P. virginiana*, respectively. Within each of the pairwise ratios, groups with different lower-case letters are significantly different based on pairwise post-hoc comparisons (*emmeans*, α = 0.05).

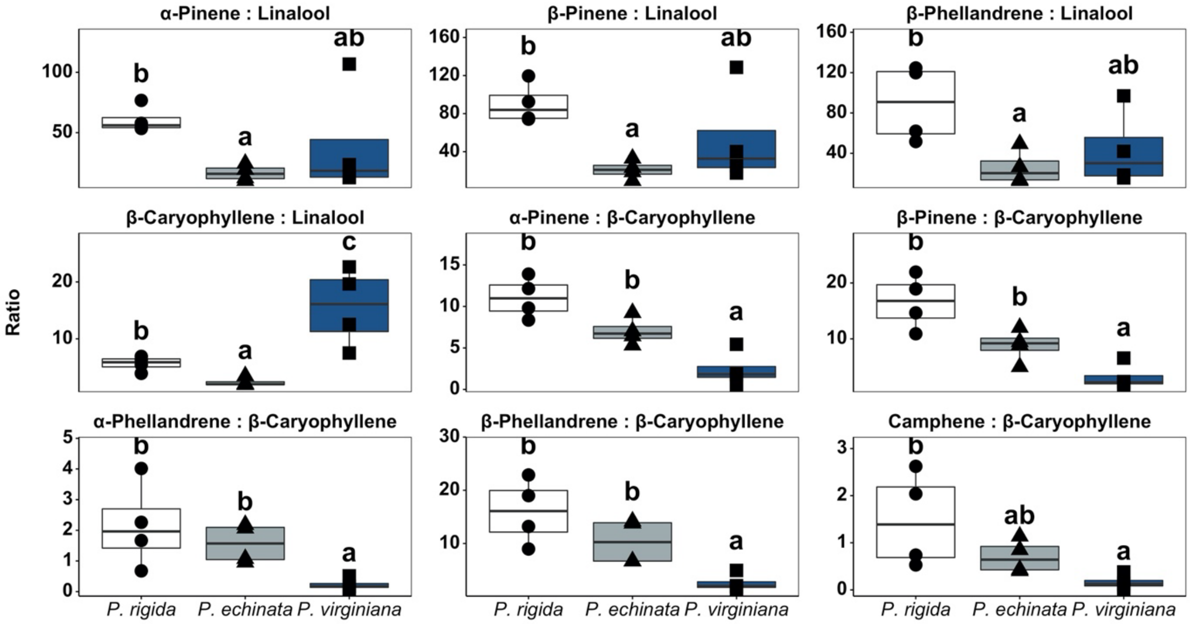

**Figure S4. C.V. error plot for ADMIXTURE runs.** Error bars represent the standard deviation around the mean C.V. error score obtained in each of 100 independent runs of ADMIXTURE for K = 1 through 10. These results indicate the optimal K is 1, which is consistent with a lack of population structure.

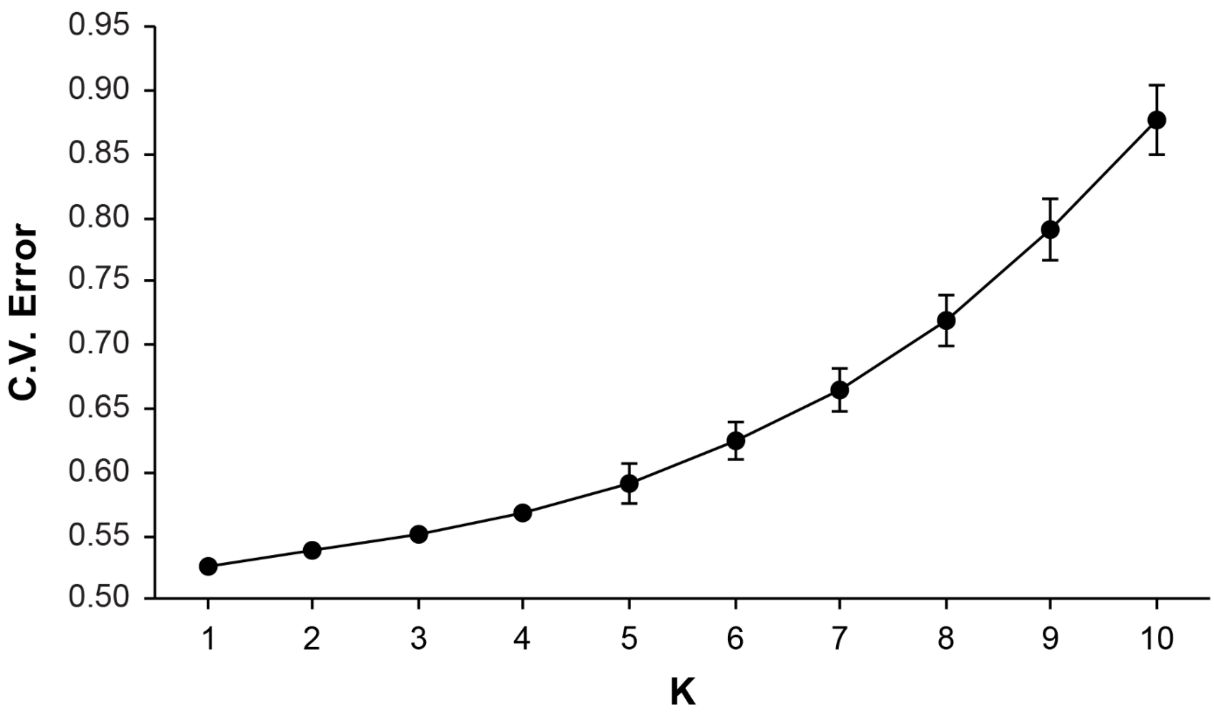

**Figure S5. Clustering solutions for K = 2.** In the chart below, each vertical bar represents one individual, and the colored bars within indicate the individual's proportion of ancestry to each of the K clusters. Dark lines segregate individuals from the Shortleaf, Pitch, and Virginia lines. There were three solutions found across the 100 runs for K = 2. The percentage of runs recovering each of the solutions below is given above the plot.

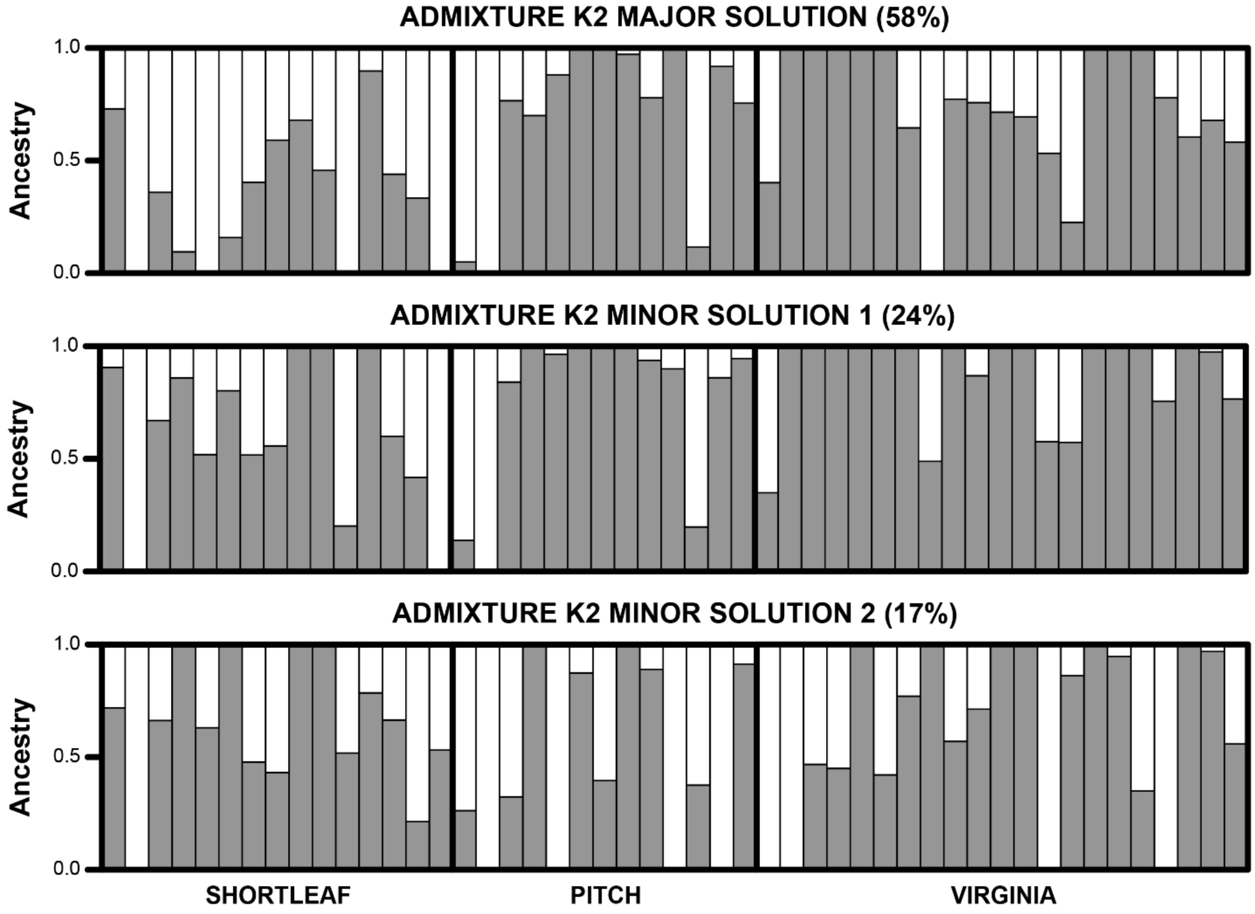

**Figure S6. Solutions for K = 3.** In the chart below, each vertical bar represents one individual, and the colored bars within indicate the individual's proportion of ancestry to each of the K clusters. Dark lines segregate individuals from the Shortleaf, Pitch, and Virginia lines. There were three solutions found across the 100 runs for K = 3. The percentage of runs recovering each of the solutions below is given above the plot.

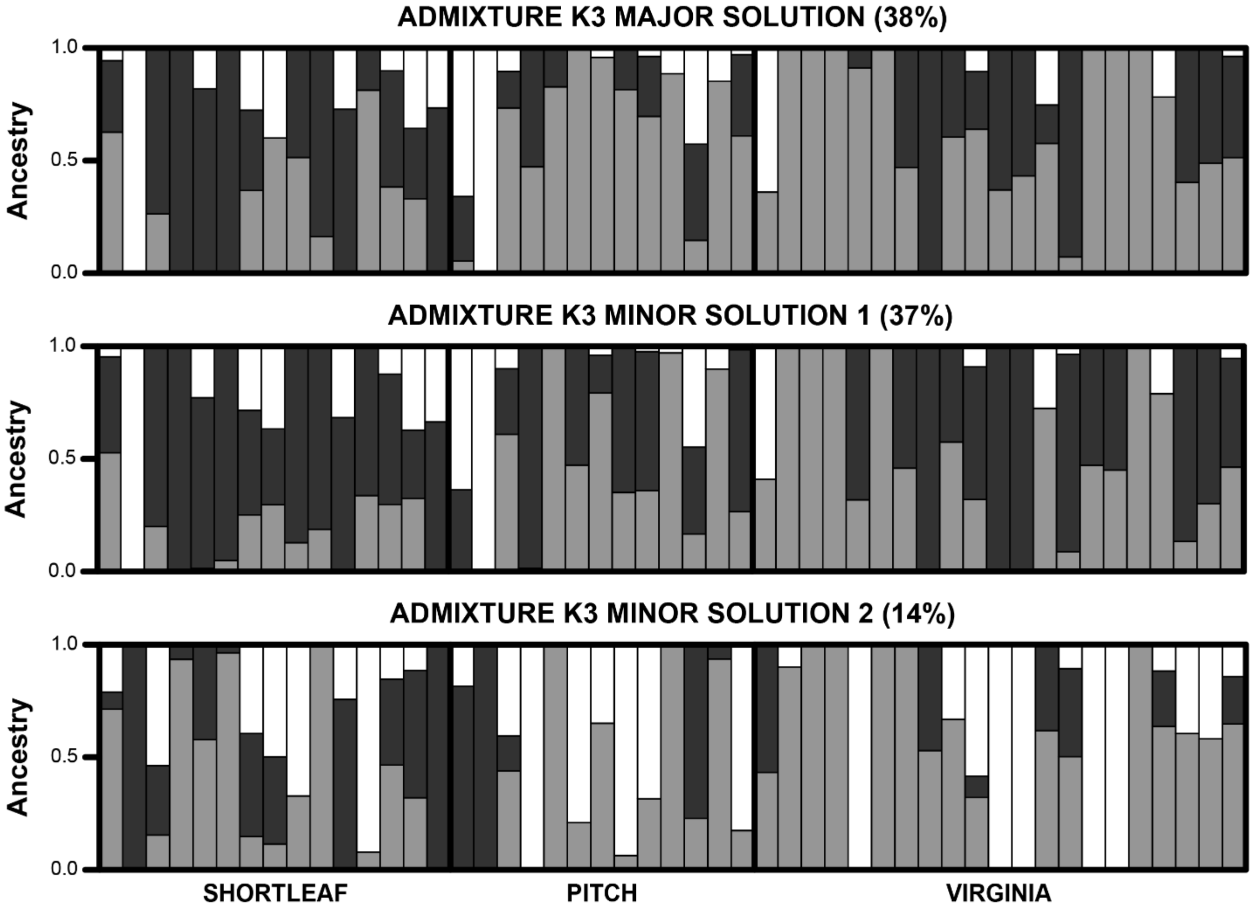

**Figure S7. Comparison of male cocoon weights.** Results for male cocoon weights across hosts were similar to those seen in females, including males from Shortleaf lines achieving the smallest cocoon weights. Males reared on *P. virginiana* were heavier than males reared on other hosts.

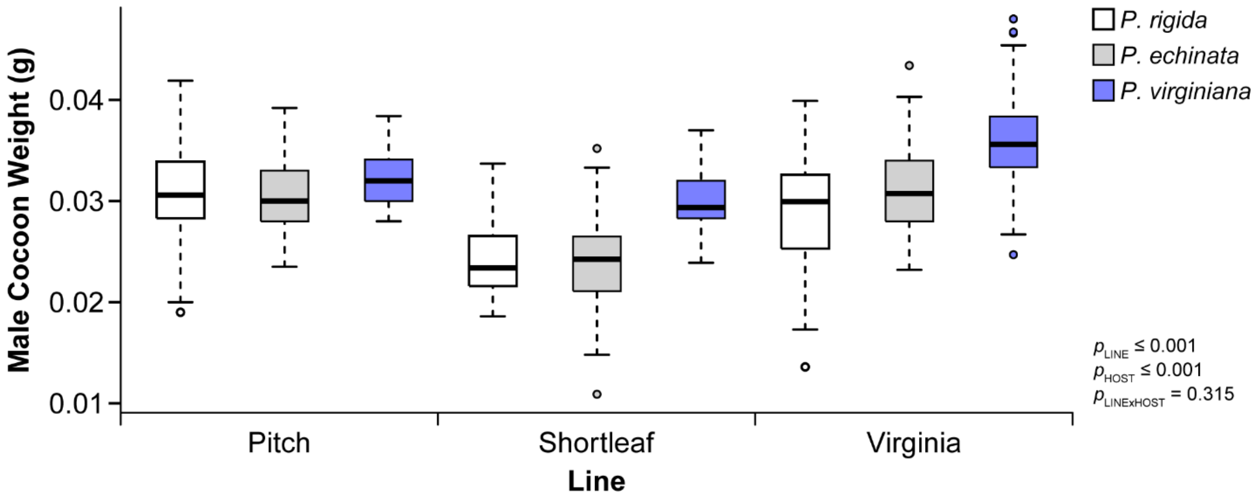
