## Supplemental Tables for "Multiple mechanisms contribute to isolation-by-environment in the redheaded pine sawfly, *Neodiprion lecontei*"

**Table S1. Collection information for resin content measurement.** Details, including site location, host tree species, collection month, and extraction details (including any base protocol deviations) for each clipping utilized in resin content measurement.

| Collection ID | Site ID | Latitude | Longitude | Tree Species | Collection Month | Filter | Hexane Purity | Ultrasonic Bath | Protocol Deviation |
|---|---|---|---|---|---|---|---|---|---|
| PS004-0 | SITE004 | 32.50772 | -83.4532 | shortleaf | May | GFF | H99 | Yes | NA |
| PS004-1 | SITE004 | 32.50772 | -83.4532 | shortleaf | May | GFD | H95 | Yes | NA |
| PS004-2 | SITE004 | 32.50772 | -83.4532 | shortleaf | May | GFD | H95 | Yes | NA |
| PS004-3 | SITE004 | 32.50772 | -83.4532 | shortleaf | May | GFD | H95 | Yes | NA |
| PS004-4 | SITE004 | 32.50772 | -83.4532 | shortleaf | May | GFD | H95 | Yes | NA |
| PS004-5 | SITE004 | 32.50772 | -83.4532 | shortleaf | May | GFD | H95 | Yes | NA |
| PS004-6 | SITE004 | 32.50772 | -83.4532 | shortleaf | May | GFD | H95 | Yes | NA |
| PS004-8 | SITE004 | 32.50772 | -83.4532 | shortleaf | May | GFF | H99 | Yes | NA |
| PS004-9 | SITE004 | 32.50772 | -83.4532 | shortleaf | May | GFD | H95 | Yes | NA |
| PS049-0 | SITE004 | 32.50772 | -83.4532 | shortleaf | August | GFD | H95 | Yes | NA |
| PS049-1 | SITE004 | 32.50772 | -83.4532 | shortleaf | August | GFD | H95 | Yes | NA |
| PS049-2 | SITE004 | 32.50772 | -83.4532 | shortleaf | August | GFD | H95 | Yes | NA |
| PS049-3 | SITE004 | 32.50772 | -83.4532 | shortleaf | August | GFD | H95 | Yes | NA |
| PS049-4 | SITE004 | 32.50772 | -83.4532 | shortleaf | August | GFD | H95 | Yes | NA |
| PS049-5 | SITE004 | 32.50772 | -83.4532 | shortleaf | August | GFD | H95 | Yes | NA |
| PS049-6 | SITE004 | 32.50772 | -83.4532 | shortleaf | August | GFD | H95 | Yes | NA |
| PS049-7 | SITE004 | 32.50772 | -83.4532 | shortleaf | August | GFD | H95 | Yes | NA |
| PS049-8 | SITE004 | 32.50772 | -83.4532 | shortleaf | August | GFD | H95 | Yes | NA |
| PS049-9 | SITE004 | 32.50772 | -83.4532 | shortleaf | August | GFD | H95 | Yes | NA |
| PS021-0 | SITE021 | 34.35316 | -87.5086 | shortleaf | May | GFD | H95 | Yes | NA |
| PS021-1 | SITE021 | 34.35316 | -87.5086 | shortleaf | May | GFD | H95 | Yes | NA |
| PS021-2 | SITE021 | 34.35316 | -87.5086 | shortleaf | May | GFD | H95 | Yes | NA |
| PS021-3 | SITE021 | 34.35316 | -87.5086 | shortleaf | May | GFD | H95 | Yes | NA |
| PS021-4 | SITE021 | 34.35316 | -87.5086 | shortleaf | May | GFD | H95 | Yes | NA |
| PS021-5 | SITE021 | 34.35316 | -87.5086 | shortleaf | May | GFD | H95 | Yes | NA |
| PS021-6 | SITE021 | 34.35316 | -87.5086 | shortleaf | May | GFD | H95 | Yes | NA |

| | | | | | | | | | |
|---|---|---|---|---|---|---|---|---|---|
| PS021-7 | SITE021 | 34.35316 | -87.5086 | shortleaf | May | GFD | H95 | Yes | NA |
| PS021-8 | SITE021 | 34.35316 | -87.5086 | shortleaf | May | GFD | H95 | Yes | NA |
| PS021-9 | SITE021 | 34.35316 | -87.5086 | shortleaf | May | GFD | H95 | Yes | NA |
| PS066-0 | SITE021 | 34.35316 | -87.5086 | shortleaf | August | GFD | H95 | No | NA |
| PS066-1 | SITE021 | 34.35316 | -87.5086 | shortleaf | August | GFD | H95 | No | NA |
| PS066-2 | SITE021 | 34.35316 | -87.5086 | shortleaf | August | GFD | H95 | No | NA |
| PS066-3 | SITE021 | 34.35316 | -87.5086 | shortleaf | August | GFD | H95 | No | NA |
| PS066-4 | SITE021 | 34.35316 | -87.5086 | shortleaf | August | GFD | H95 | No | NA |
| PS066-5 | SITE021 | 34.35316 | -87.5086 | shortleaf | August | GFD | H95 | No | NA |
| PS066-6 | SITE021 | 34.35316 | -87.5086 | shortleaf | August | GFD | H95 | No | NA |
| PS066-7 | SITE021 | 34.35316 | -87.5086 | shortleaf | August | GFD | H95 | No | NA |
| PS066-8 | SITE021 | 34.35316 | -87.5086 | shortleaf | August | GFD | H95 | No | NA |
| PS066-9 | SITE021 | 34.35316 | -87.5086 | shortleaf | August | GFD | H95 | No | NA |
| PS023-0 | SITE023 | 35.94187 | -86.5271 | virginia | May | GFD | H95 | Yes | NA |
| PS023-1 | SITE023 | 35.94187 | -86.5271 | virginia | May | GFD | H95 | Yes | NA |
| PS023-2 | SITE023 | 35.94187 | -86.5271 | virginia | May | GFD | H95 | Yes | NA |
| PS023-3 | SITE023 | 35.94187 | -86.5271 | virginia | May | GFD | H95 | Yes | NA |
| PS023-4 | SITE023 | 35.94187 | -86.5271 | virginia | May | GFD | H95 | Yes | NA |
| PS023-5 | SITE023 | 35.94187 | -86.5271 | virginia | May | GFD | H95 | Yes | NA |
| PS023-7 | SITE023 | 35.94187 | -86.5271 | virginia | May | GFD | H95 | Yes | NA |
| PS023-8 | SITE023 | 35.94187 | -86.5271 | virginia | May | GFD | H95 | Yes | NA |
| PS068-0 | SITE023 | 35.94187 | -86.5271 | virginia | August | GFD | H95 | Yes | NA |
| PS068-1 | SITE023 | 35.94187 | -86.5271 | virginia | August | GFD | H95 | Yes | NA |
| PS068-2 | SITE023 | 35.94187 | -86.5271 | virginia | August | GFD | H95 | Yes | NA |
| PS068-3 | SITE023 | 35.94187 | -86.5271 | virginia | August | GFD | H95 | Yes | NA |
| PS068-4 | SITE023 | 35.94187 | -86.5271 | virginia | August | GFD | H95 | Yes | NA |
| PS068-5 | SITE023 | 35.94187 | -86.5271 | virginia | August | GFD | H95 | Yes | NA |
| PS068-6 | SITE023 | 35.94187 | -86.5271 | virginia | August | GFD | H95 | Yes | NA |
| PS068-7 | SITE023 | 35.94187 | -86.5271 | virginia | August | GFD | H95 | Yes | NA |
| PS068-8 | SITE023 | 35.94187 | -86.5271 | virginia | August | GFD | H95 | Yes | NA |
| PS068-9 | SITE023 | 35.94187 | -86.5271 | virginia | August | GFD | H95 | Yes | NA |

| | | | | | | | | | |
|---|---|---|---|---|---|---|---|---|---|
| PS024-0 | SITE024 | 38.16739 | -83.5905 | virginia | May | GFD | H95 | Yes | NA |
| PS024-1 | SITE024 | 38.16739 | -83.5905 | virginia | May | GFD | H95 | Yes | NA |
| PS024-2 | SITE024 | 38.16739 | -83.5905 | virginia | May | GFD | H95 | Yes | NA |
| PS024-3 | SITE024 | 38.16739 | -83.5905 | virginia | May | GFD | H95 | Yes | NA |
| PS024-4 | SITE024 | 38.16739 | -83.5905 | virginia | May | GFD | H95 | Yes | NA |
| PS024-5 | SITE024 | 38.16739 | -83.5905 | virginia | May | GFD | H95 | Yes | NA |
| PS024-6 | SITE024 | 38.16739 | -83.5905 | virginia | May | GFD | H95 | Yes | NA |
| PS024-7 | SITE024 | 38.16739 | -83.5905 | virginia | May | GFD | H95 | Yes | NA |
| PS024-8 | SITE024 | 38.16739 | -83.5905 | virginia | May | GFD | H95 | Yes | NA |
| PS024-9 | SITE024 | 38.16739 | -83.5905 | virginia | May | GFD | H95 | Yes | NA |
| PS070-0 | SITE024 | 38.16739 | -83.5905 | virginia | August | GFD | H95 | Yes | NA |
| PS070-1 | SITE024 | 38.16739 | -83.5905 | virginia | August | GFD | H95 | Yes | NA |
| PS070-2 | SITE024 | 38.16739 | -83.5905 | virginia | August | GFD | H95 | Yes | NA |
| PS070-3 | SITE024 | 38.16739 | -83.5905 | virginia | August | GFD | H95 | Yes | NA |
| PS070-4 | SITE024 | 38.16739 | -83.5905 | virginia | August | GFD | H95 | Yes | NA |
| PS070-5 | SITE024 | 38.16739 | -83.5905 | virginia | August | GFD | H95 | Yes | NA |
| PS070-6 | SITE024 | 38.16739 | -83.5905 | virginia | August | GFD | H95 | Yes | NA |
| PS070-7 | SITE024 | 38.16739 | -83.5905 | virginia | August | GFD | H95 | Yes | NA |
| PS070-8 | SITE024 | 38.16739 | -83.5905 | virginia | August | GFD | H95 | Yes | NA |
| PS070-9 | SITE024 | 38.16739 | -83.5905 | virginia | August | GFD | H95 | Yes | NA |
| PS026-0 | SITE026 | 35.47462 | -80.262 | shortleaf | May | GFD | H95 | Yes | NA |
| PS026-1 | SITE026 | 35.47462 | -80.262 | shortleaf | May | GFD | H95 | Yes | NA |
| PS026-2 | SITE026 | 35.47462 | -80.262 | shortleaf | May | GFD | H95 | Yes | NA |
| PS026-3 | SITE026 | 35.47462 | -80.262 | shortleaf | May | GFD | H95 | Yes | NA |
| PS026-4 | SITE026 | 35.47462 | -80.262 | shortleaf | May | GFD | H95 | Yes | NA |
| PS026-5 | SITE026 | 35.47462 | -80.262 | shortleaf | May | GFD | H95 | Yes | NA |
| PS026-6 | SITE026 | 35.47462 | -80.262 | shortleaf | May | GFD | H95 | Yes | NA |
| PS026-7 | SITE026 | 35.47462 | -80.262 | shortleaf | May | GFD | H95 | Yes | NA |
| PS026-8 | SITE026 | 35.47462 | -80.262 | shortleaf | May | GFD | H95 | Yes | NA |
| PS026-9 | SITE026 | 35.47462 | -80.262 | shortleaf | May | GFD | H95 | Yes | NA |

| | | | | | | | | | |
|---|---|---|---|---|---|---|---|---|---|
| PS072-0 | SITE026 | 35.47462 | -80.262 | shortleaf | August | GFD | H95 | No | First Transfer 1 Day Late; Second Transfer Early; Third Transfer Late |
| PS072-1 | SITE026 | 35.47462 | -80.262 | shortleaf | August | GFD | H95 | No | First Transfer 1 Day Late; Second Transfer Early; Third Transfer Late |
| PS072-2 | SITE026 | 35.47462 | -80.262 | shortleaf | August | GFD | H95 | No | First Transfer 1 Day Late; Second Transfer Early; Third Transfer Late |
| PS072-3 | SITE026 | 35.47462 | -80.262 | shortleaf | August | GFD | H95 | No | First Transfer 1 Day Late; Second Transfer Early; Third Transfer Late |
| PS072-4 | SITE026 | 35.47462 | -80.262 | shortleaf | August | GFD | H95 | No | First Transfer 1 Day Late; Second Transfer Early; Third Transfer Late |
| PS072-5 | SITE026 | 35.47462 | -80.262 | shortleaf | August | GFD | H95 | No | First Transfer 1 Day Late; Second Transfer Early; Third Transfer Late |
| PS072-6 | SITE026 | 35.47462 | -80.262 | shortleaf | August | GFD | H95 | No | First Transfer 1 Day Late; Second Transfer Early; Third Transfer Late |
| PS072-7 | SITE026 | 35.47462 | -80.262 | shortleaf | August | GFD | H95 | No | First Transfer 1 Day Late; Second Transfer Early; Third Transfer Late |
| PS072-8 | SITE026 | 35.47462 | -80.262 | shortleaf | August | GFD | H95 | No | First Transfer 1 Day Late; Second Transfer Early; Third Transfer Late |
| PS072-9 | SITE026 | 35.47462 | -80.262 | shortleaf | August | GFD | H95 | No | First Transfer 1 Day Late; Second Transfer Early; Third Transfer Late |
| PS028-0 | SITE028 | 37.11326 | -78.0271 | virginia | May | GFD | H95 | Yes | NA |
| PS028-1 | SITE028 | 37.11326 | -78.0271 | virginia | May | GFD | H95 | Yes | NA |
| PS028-2 | SITE028 | 37.11326 | -78.0271 | virginia | May | GFD | H95 | Yes | NA |
| PS028-3 | SITE028 | 37.11326 | -78.0271 | virginia | May | GFD | H95 | Yes | NA |
| PS028-4 | SITE028 | 37.11326 | -78.0271 | virginia | May | GFD | H95 | Yes | NA |
| PS028-5 | SITE028 | 37.11326 | -78.0271 | virginia | May | GFD | H95 | Yes | NA |

| | | | | | | | | | |
|---|---|---|---|---|---|---|---|---|---|
| PS028-6 | SITE028 | 37.11326 | -78.0271 | virginia | May | GFD | H95 | Yes | NA |
| PS028-7 | SITE028 | 37.11326 | -78.0271 | virginia | May | GFD | H95 | Yes | NA |
| PS028-8 | SITE028 | 37.11326 | -78.0271 | virginia | May | GFD | H95 | Yes | NA |
| PS028-9 | SITE028 | 37.11326 | -78.0271 | virginia | May | GFD | H95 | Yes | NA |
| PS074-0 | SITE028 | 37.11326 | -78.0271 | virginia | August | GFD | H95 | Yes | NA |
| PS074-1 | SITE028 | 37.11326 | -78.0271 | virginia | August | GFD | H95 | Yes | NA |
| PS074-2 | SITE028 | 37.11326 | -78.0271 | virginia | August | GFD | H95 | Yes | NA |
| PS074-3 | SITE028 | 37.11326 | -78.0271 | virginia | August | GFD | H95 | Yes | NA |
| PS074-4 | SITE028 | 37.11326 | -78.0271 | virginia | August | GFD | H95 | Yes | NA |
| PS074-5 | SITE028 | 37.11326 | -78.0271 | virginia | August | GFD | H95 | Yes | NA |
| PS074-6 | SITE028 | 37.11326 | -78.0271 | virginia | August | GFD | H95 | Yes | NA |
| PS074-7 | SITE028 | 37.11326 | -78.0271 | virginia | August | GFD | H95 | Yes | NA |
| PS074-8 | SITE028 | 37.11326 | -78.0271 | virginia | August | GFD | H95 | Yes | NA |
| PS074-9 | SITE028 | 37.11326 | -78.0271 | virginia | August | GFD | H95 | Yes | NA |
| PS029-0 | SITE029 | 38.72832 | -79.4632 | pitch | May | GFD | H95 | Yes | NA |
| PS029-1 | SITE029 | 38.72832 | -79.4632 | pitch | May | GFD | H95 | Yes | NA |
| PS029-2 | SITE029 | 38.72832 | -79.4632 | pitch | May | GFD | H95 | Yes | NA |
| PS029-3 | SITE029 | 38.72832 | -79.4632 | pitch | May | GFD | H95 | Yes | NA |
| PS029-4 | SITE029 | 38.72832 | -79.4632 | pitch | May | GFD | H95 | Yes | NA |
| PS029-5 | SITE029 | 38.72832 | -79.4632 | pitch | May | GFD | H95 | Yes | NA |
| PS029-6 | SITE029 | 38.72832 | -79.4632 | pitch | May | GFD | H95 | Yes | NA |
| PS029-7 | SITE029 | 38.72832 | -79.4632 | pitch | May | GFD | H95 | Yes | NA |
| PS029-8 | SITE029 | 38.72832 | -79.4632 | pitch | May | GFD | H95 | Yes | NA |
| PS075-0 | SITE029 | 38.72832 | -79.4632 | pitch | August | GFD | H95 | Yes | NA |
| PS075-1 | SITE029 | 38.72832 | -79.4632 | pitch | August | GFD | H95 | Yes | NA |
| PS075-2 | SITE029 | 38.72832 | -79.4632 | pitch | August | GFD | H95 | Yes | NA |
| PS075-3 | SITE029 | 38.72832 | -79.4632 | pitch | August | GFD | H95 | Yes | NA |
| PS075-4 | SITE029 | 38.72832 | -79.4632 | pitch | August | GFD | H95 | Yes | NA |
| PS075-5 | SITE029 | 38.72832 | -79.4632 | pitch | August | GFD | H95 | Yes | NA |
| PS075-6 | SITE029 | 38.72832 | -79.4632 | pitch | August | GFD | H95 | Yes | NA |
| PS075-7 | SITE029 | 38.72832 | -79.4632 | pitch | August | GFD | H95 | Yes | NA |

| | | | | | | | | | |
|---|---|---|---|---|---|---|---|---|---|
| PS075-8 | SITE029 | 38.72832 | -79.4632 | pitch | August | GFD | H95 | Yes | NA |
| PS075-9 | SITE029 | 38.72832 | -79.4632 | pitch | August | GFD | H95 | Yes | NA |
| PS030-0 | SITE030 | 38.72832 | -79.4632 | virginia | May | GFD | H95 | Yes | NA |
| PS030-1 | SITE030 | 38.72832 | -79.4632 | virginia | May | GFD | H95 | Yes | NA |
| PS030-2 | SITE030 | 38.72832 | -79.4632 | virginia | May | GFD | H95 | Yes | NA |
| PS030-3 | SITE030 | 38.72832 | -79.4632 | virginia | May | GFD | H95 | Yes | NA |
| PS030-4 | SITE030 | 38.72832 | -79.4632 | virginia | May | GFD | H95 | Yes | NA |
| PS030-5 | SITE030 | 38.72832 | -79.4632 | virginia | May | GFD | H95 | Yes | NA |
| PS030-7 | SITE030 | 38.72832 | -79.4632 | virginia | May | GFD | H95 | Yes | NA |
| PS030-8 | SITE030 | 38.72832 | -79.4632 | virginia | May | GFD | H95 | Yes | NA |
| PS076-0 | SITE030 | 38.72832 | -79.4632 | virginia | August | GFD | H95 | Yes | NA |
| PS076-1 | SITE030 | 38.72832 | -79.4632 | virginia | August | GFD | H95 | Yes | NA |
| PS076-2 | SITE030 | 38.72832 | -79.4632 | virginia | August | GFD | H95 | Yes | NA |
| PS076-3 | SITE030 | 38.72832 | -79.4632 | virginia | August | GFD | H95 | Yes | NA |
| PS076-4 | SITE030 | 38.72832 | -79.4632 | virginia | August | GFD | H95 | Yes | NA |
| PS076-5 | SITE030 | 38.72832 | -79.4632 | virginia | August | GFD | H95 | Yes | NA |
| PS076-6 | SITE030 | 38.72832 | -79.4632 | virginia | August | GFD | H95 | Yes | NA |
| PS076-7 | SITE030 | 38.72832 | -79.4632 | virginia | August | GFD | H95 | Yes | NA |
| PS076-8 | SITE030 | 38.72832 | -79.4632 | virginia | August | GFD | H95 | Yes | NA |
| PS076-9 | SITE030 | 38.72832 | -79.4632 | virginia | August | GFD | H95 | Yes | NA |
| PS032-0 | SITE032 | 39.88555 | -74.5062 | pitch | May | GFD | H95 | Yes | NA |
| PS032-3 | SITE032 | 39.88555 | -74.5062 | pitch | May | GFD | H95 | Yes | NA |
| PS032-4 | SITE032 | 39.88555 | -74.5062 | pitch | May | GFD | H95 | Yes | NA |
| PS032-5 | SITE032 | 39.88555 | -74.5062 | pitch | May | GFD | H95 | Yes | NA |
| PS032-6 | SITE032 | 39.88555 | -74.5062 | pitch | May | GFD | H95 | Yes | NA |
| PS032-7 | SITE032 | 39.88555 | -74.5062 | pitch | May | GFD | H95 | Yes | NA |
| PS032-8 | SITE032 | 39.88555 | -74.5062 | pitch | May | GFD | H95 | Yes | NA |
| PS032-9 | SITE032 | 39.88555 | -74.5062 | pitch | May | GFD | H95 | Yes | NA |
| PS078-0 | SITE032 | 39.88555 | -74.5062 | pitch | August | GFD | H95 | Yes | NA |
| PS078-1 | SITE032 | 39.88555 | -74.5062 | pitch | August | GFD | H95 | Yes | NA |
| PS078-2 | SITE032 | 39.88555 | -74.5062 | pitch | August | GFD | H95 | Yes | NA |

| | | | | | | | | | |
|---|---|---|---|---|---|---|---|---|---|
| PS078-3 | SITE032 | 39.88555 | -74.5062 | pitch | August | GFD | H95 | Yes | NA |
| PS078-4 | SITE032 | 39.88555 | -74.5062 | pitch | August | GFD | H95 | Yes | NA |
| PS078-5 | SITE032 | 39.88555 | -74.5062 | pitch | August | GFD | H95 | Yes | NA |
| PS078-6 | SITE032 | 39.88555 | -74.5062 | pitch | August | GFD | H95 | Yes | NA |
| PS078-7 | SITE032 | 39.88555 | -74.5062 | pitch | August | GFD | H95 | Yes | NA |
| PS078-8 | SITE032 | 39.88555 | -74.5062 | pitch | August | GFD | H95 | Yes | NA |
| PS078-9 | SITE032 | 39.88555 | -74.5062 | pitch | August | GFD | H95 | Yes | NA |
| PS033-0 | SITE033 | 41.8465 | -70.6793 | pitch | May | GFD | H95 | Yes | NA |
| PS033-1 | SITE033 | 41.8465 | -70.6793 | pitch | May | GFD | H95 | Yes | NA |
| PS033-2 | SITE033 | 41.8465 | -70.6793 | pitch | May | GFD | H95 | Yes | NA |
| PS033-3 | SITE033 | 41.8465 | -70.6793 | pitch | May | GFD | H95 | Yes | NA |
| PS033-4 | SITE033 | 41.8465 | -70.6793 | pitch | May | GFD | H95 | Yes | NA |
| PS033-5 | SITE033 | 41.8465 | -70.6793 | pitch | May | GFD | H95 | Yes | NA |
| PS033-6 | SITE033 | 41.8465 | -70.6793 | pitch | May | GFD | H95 | Yes | NA |
| PS033-7 | SITE033 | 41.8465 | -70.6793 | pitch | May | GFD | H95 | Yes | NA |
| PS033-8 | SITE033 | 41.8465 | -70.6793 | pitch | May | GFD | H95 | Yes | NA |
| PS033-9 | SITE033 | 41.8465 | -70.6793 | pitch | May | GFD | H95 | Yes | NA |
| PS079-0 | SITE033 | 41.8465 | -70.6793 | pitch | August | GFD | H95 | No | First Transfer 1 Day Late; Second Transfer Early; Third Transfer Late |
| PS079-1 | SITE033 | 41.8465 | -70.6793 | pitch | August | GFD | H95 | No | First Transfer 1 Day Late; Second Transfer Early; Third Transfer Late |
| PS079-2 | SITE033 | 41.8465 | -70.6793 | pitch | August | GFD | H95 | No | First Transfer 1 Day Late; Second Transfer Early; Third Transfer Late |
| PS079-3 | SITE033 | 41.8465 | -70.6793 | pitch | August | GFD | H95 | No | First Transfer 1 Day Late; Second Transfer Early; Third Transfer Late |
| PS079-4 | SITE033 | 41.8465 | -70.6793 | pitch | August | GFD | H95 | No | First Transfer 1 Day Late; Second Transfer Early; Third Transfer Late |

| | | | | | | | | | |
|---|---|---|---|---|---|---|---|---|---|
| PS079-5 | SITE033 | 41.8465 | -70.6793 | pitch | August | GFD | H95 | No | First Transfer 1 Day Late; Second Transfer Early; Third Transfer Late |
| PS079-6 | SITE033 | 41.8465 | -70.6793 | pitch | August | GFD | H95 | No | First Transfer 1 Day Late; Second Transfer Early; Third Transfer Late |
| PS079-7 | SITE033 | 41.8465 | -70.6793 | pitch | August | GFD | H95 | No | First Transfer 1 Day Late; Second Transfer Early; Third Transfer Late |
| PS079-8 | SITE033 | 41.8465 | -70.6793 | pitch | August | GFD | H95 | No | First Transfer 1 Day Late; Second Transfer Early; Third Transfer Late |
| PS079-9 | SITE033 | 41.8465 | -70.6793 | pitch | August | GFD | H95 | No | First Transfer 1 Day Late; Second Transfer Early; Third Transfer Late |
| PS045-0 | SITE045 | 37.26224 | -84.9634 | pitch | May | GFD | H95 | Yes | NA |
| PS045-1 | SITE045 | 37.26224 | -84.9634 | pitch | May | GFD | H95 | Yes | NA |
| PS045-2 | SITE045 | 37.26224 | -84.9634 | pitch | May | GFD | H95 | Yes | NA |
| PS045-3 | SITE045 | 37.26224 | -84.9634 | pitch | May | GFD | H95 | Yes | NA |
| PS045-4 | SITE045 | 37.26224 | -84.9634 | pitch | May | GFD | H95 | Yes | NA |
| PS045-5 | SITE045 | 37.26224 | -84.9634 | pitch | May | GFD | H95 | Yes | NA |
| PS045-6 | SITE045 | 37.26224 | -84.9634 | pitch | May | GFD | H95 | Yes | NA |
| PS045-7 | SITE045 | 37.26224 | -84.9634 | pitch | May | GFD | H95 | Yes | NA |
| PS045-8 | SITE045 | 37.26224 | -84.9634 | pitch | May | GFD | H95 | Yes | NA |
| PS045-9 | SITE045 | 37.26224 | -84.9634 | pitch | May | GFD | H95 | Yes | NA |
| PS069-0 | SITE045 | 37.26224 | -84.9634 | pitch | August | GFD | H95 | Yes | NA |
| PS069-1 | SITE045 | 37.26224 | -84.9634 | pitch | August | GFD | H95 | Yes | NA |
| PS069-2 | SITE045 | 37.26224 | -84.9634 | pitch | August | GFD | H95 | Yes | NA |
| PS069-3 | SITE045 | 37.26224 | -84.9634 | pitch | August | GFD | H95 | Yes | NA |
| PS069-4 | SITE045 | 37.26224 | -84.9634 | pitch | August | GFD | H95 | Yes | NA |
| PS069-5 | SITE045 | 37.26224 | -84.9634 | pitch | August | GFD | H95 | Yes | NA |
| PS069-6 | SITE045 | 37.26224 | -84.9634 | pitch | August | GFD | H95 | Yes | NA |
| PS069-7 | SITE045 | 37.26224 | -84.9634 | pitch | August | GFD | H95 | Yes | NA |

| | | | | | | | | | |
|---|---|---|---|---|---|---|---|---|---|
| PS069-8 | SITE045 | 37.26224 | -84.9634 | pitch | August | GFD | H95 | Yes | NA |
| PS069-9 | SITE045 | 37.26224 | -84.9634 | pitch | August | GFD | H95 | Yes | NA |

**Table S2. Usage for each *Neodiprion lecontei* colony.** The collection date and source host plant colonies are given for each colony. An "x" denotes if that colony was used for genetic work (RAD Sequencing), analysis of ovipositor shape (Ovipositor Morphology), IBE mechanism analyses (Ecological Assays), and/or assessments of phenological differences (Temporal Isolation).

| ID | Collection Date | Source Host Plant | RAD Sequencing | Ovipositor Morphology | Ecological Assays | Temporal Isolation |
|---|---|---|---|---|---|---|
| LL002 | 6/13/2012 | *P. echinata* | x | | | |
| LL003 | 6/13/2012 | *P. echinata* | x | | | |
| LL004 | 6/13/2012 | *P. echinata* | x | | | |
| LL005 | 6/15/2012 | *P. rigida* | x | | | |
| LL006 | 6/15/2012 | *P. rigida* | x | | | |
| LL027 | 7/25/2013 | *P. echinata* | x | | | |
| LL028 | 8/2/2013 | *P. echinata* | x | | | x |
| LL047 | 6/9/2014 | *P. echinata* | | x | | |
| LL049 | 6/9/2014 | *P. echinata* | | x | | |
| LL051 | 6/9/2014 | *P. echinata* | | x | | |
| LL053 | 6/9/2014 | *P. virginiana* | x | | | |
| LL054 | 6/9/2014 | *P. virginiana* | | x | | x |
| LL055 | 6/9/2014 | *P. virginiana* | | | | x |
| LL058 | 6/19/2014 | *P. virginiana* | x | x | | x |
| LL059 | 6/19/2014 | *P. virginiana* | | x | | x |
| LL061 | 6/19/2014 | *P. virginiana* | x | x | | x |
| LL062 | 6/19/2014 | *P. echinata* | x | x | | x |
| LL063 | 6/19/2014 | *P. echinata* | | | | x |
| LL064 | 6/19/2014 | *P. rigida* | x | | | |
| LL066 | 6/19/2014 | *P. rigida* | | x | | x |
| LL067 | 6/19/2014 | *P. rigida* | | x | | x |
| LL069 | 6/27/2014 | *P. virginiana* | x | | | |
| LL070 | 6/27/2014 | *P. rigida* | x | x | | |
| LL071 | 6/27/2014 | *P. echinata* | x | | | |
| LL072 | 6/27/2014 | *P. virginiana* | x | | | |
| LL073 | 6/27/2014 | *P. echinata* | x | | | |
| LL074 | 6/30/2014 | *P. echinata* | x | x | | x |
| LL075 | 6/30/2014 | *P. rigida* | | x | | x |
| LL076 | 6/30/2014 | *P. virginiana* | | x | | x |
| LL078 | 6/30/2014 | *P. rigida* | x | | | |
| LL092 | 8/28/2014 | *P. virginiana* | x | | | |
| LL102 | 9/4/2014 | *P. rigida* | x | | | |
| LL116 | 6/2/2015 | *P. rigida* | | x | | |
| LL121 | 6/17/2015 | *P. rigida* | | x | | |
| LL122 | 6/17/2015 | *P. rigida* | | x | | |

| | | | | | | |
|---|---|---|---|---|---|---|
| LL136 | 6/19/2015 | *P. virginiana* | x | | | |
| LL137 | 6/19/2015 | *P. virginiana* | x | | | |
| LL139 | 6/19/2015 | *P. echinata* | x | | | |
| LL140 | 6/19/2015 | *P. virginiana* | x | | | |
| LL179 | 7/3/2015 | *P. echinata* | x | | | |
| LL183 | 7/8/2015 | *P. echinata* | | x | | |
| LL190 | 7/14/2015 | *P. echinata* | | x | | |
| LL191 | 7/20/2015 | *P. echinata* | | x | | |
| LL216 | 8/28/2015 | *P. echinata* | | x | | |
| RB076 | 7/5/2012 | *P. virginiana* | x | | | |
| RB122 | 8/20/2012 | *P. echinata* | x | | | |
| RB126 | 8/31/2012 | *P. virginiana* | x | | | |
| RB127 | 8/31/2012 | *P. virginiana* | x | | | |
| RB128 | 8/31/2012 | *P. virginiana* | x | | | |
| RB129 | 8/31/2012 | *P. echinata* | x | | | |
| RB132 | 9/14/2012 | *P. rigida* | x | | | |
| RB135 | 9/14/2012 | *P. virginiana* | x | | | |
| RB335 | 8/22/2013 | *P. echinata* | | | x | x |
| RB336 | 8/22/2013 | *P. echinata* | | | x | x |
| RB337 | 8/22/2013 | *P. virginiana* | x | | x | x |
| RB338 | 8/22/2013 | *P. echinata* | | | x | |
| RB339 | 8/22/2013 | *P. echinata* | x | | x | x |
| RB341 | 8/22/2013 | *P. virginiana* | | | x | x |
| RB342 | 9/9/2013 | *P. echinata* | | | x | x |
| RB343 | 9/9/2013 | *P. rigida* | x | | x | x |
| RB344 | 9/9/2013 | *P. rigida* | x | | x | x |
| RB345 | 9/9/2013 | *P. rigida* | | | x | x |
| RB346 | 9/9/2013 | *P. virginiana* | | | x | x |
| RB347 | 9/9/2013 | *P. virginiana* | x | | x | x |
| RB348 | 9/9/2013 | *P. virginiana* | | | x | |
| RB349 | 9/9/2013 | *P. echinata* | x | | | |
| RB353 | 9/9/2013 | *P. echinata* | x | | | |
| RB354 | 9/9/2013 | *P. virginiana* | | | x | x |
| RB355 | 9/9/2013 | *P. virginiana* | x | | | |
| RB357 | 9/9/2013 | *P. virginiana* | x | | | |
| RB358 | 9/9/2013 | *P. virginiana* | x | | | |

**Table S3. Sequencing information for each specimen.** The PCR index ("Library") and unique, variable-length barcode ("Barcode") for each individual is given. Various sequencing statistics, including base read numbers ("Raw Reads"), reads surviving quality filters ("Filtered Reads"), uniquely mapping, non-PCR duplicate alignments ("Retained Alignments"), RAD loci formed in STACKS ("RAD Loci"), average coverage within formed RAD loci ("Mean Coverage"), percent missing data ("Missing Data"), and the proportion of heterozygous sites, which was used to identify putative haploid individuals ("Proportion Heterozygous Sites"). Putative haploid males are marked with the male symbol (♂) and confirmed adult females are marked with the female symbol (♀). All other specimens are diploid larvae of unknown sex.

| ID | Library | Barcode | Line | Raw Reads | Filtered Reads | Retained Alignments | RAD Loci | Mean Coverage | Missing Data | Proportion Heterozygous Sites |
|---|---|---|---|---|---|---|---|---|---|---|
| LL002_01 | GTGAAACG | GAGCGACAT | Shortleaf | 1557242 | 1547025 | 862149 | 18561 | 42 | 0.42 | 0.2887 |
| LL003_01 ♂ | GTGAAACG | CCTTGCCATT | Shortleaf | 845863 | 844862 | 490193 | 15677 | 27 | 5.13 | 0.0033 |
| LL004_01♀ | ATCACGAT | TGGCACAGA | Shortleaf | 3392337 | 3386292 | 1760807 | 25507 | 64 | 1.11 | 0.1657 |
| LL005_01♀ | TGACCAAT | TGGCAACAGA | Pitch | 2859217 | 2854217 | 1464252 | 20530 | 64 | 0.64 | 0.2839 |
| LL006_02b | GTGAAACG | TCAGAGAT | Pitch | 1075134 | 1073489 | 604364 | 17004 | 32 | 1.67 | 0.2826 |
| LL027 | CGATGTAT | TGGCAACAGA | Shortleaf | 12597 | 12568 | 7738 | 1 | 25 | 100 | N/A |
| LL028 | TTAGGCAT | CTCTCGCAT | Shortleaf | 17161130 | 17135652 | 6182743 | 37787 | 153 | 0.74 | 0.1797 |
| LL053 | TGACCAAT | TCTTGG | Virginia | 1973784 | 1955963 | 1043020 | 18657 | 50 | 0.92 | 0.2835 |
| LL058_1 | TTAGGCAT | GAAGTG | Virginia | 373279 | 367092 | 213856 | 8805 | 17 | 42.57 | 0.2565 |
| LL058.2 | AGTCAACA | ACGGTACT | Virginia | 1638435 | 1635814 | 873260 | 18222 | 44 | 3.00 | 0.2971 |
| LL058_3R | ATCACGAT | GCAAGCCAT | Virginia | 2781563 | 2777303 | 1491987 | 22967 | 58 | 0.20 | 0.2625 |
| LL061 | TGACCAAT | ACCAGGA | Virginia | 2193648 | 2190005 | 1166563 | 19059 | 55 | 0.56 | 0.2736 |
| LL062_1 | CGATGTAT | TGACGCCA | Shortleaf | 595932 | 594914 | 340064 | 10774 | 27 | 29.21 | 0.2269 |
| LL062_2 | TGACCAAT | GAGCGACAT | Shortleaf | 2863779 | 2843518 | 1477165 | 20257 | 66 | 0.32 | 0.2646 |
| LL062_3R | GTCCGCAC | AACGTGCCT | Shortleaf | 2402042 | 2398243 | 1279398 | 19166 | 61 | 0.45 | 0.2758 |
| LL064 | TTAGGCAT | ACGGTACT | Pitch | 2945519 | 2940999 | 1509853 | 19472 | 70 | 0.42 | 0.2544 |
| LL069_1R | TTAGGCAT | TGCTT | Virginia | 860914 | 849898 | 468081 | 15875 | 26 | 4.85 | 0.2517 |
| LL069_2R | ATCACGAT | TGACGCCA | Virginia | 1756970 | 1754231 | 953974 | 19938 | 42 | 0.29 | 0.2706 |
| LL069_3 | GTCCGCAC | GGAACGA | Virginia | 1584987 | 1582254 | 858881 | 17292 | 44 | 0.69 | 0.2229 |
| LL070_1 | CGATGTAT | ATATCGCCA | Pitch | 536460 | 535612 | 294868 | 10129 | 24 | 33.08 | 0.2594 |
| LL070_2 | TTAGGCAT | AACGCACATT | Pitch | 1874819 | 1871930 | 1051433 | 19154 | 50 | 2.53 | 0.2459 |
| LL070_3R | CGATGTAT | AACGTGCCT | Pitch | 1755110 | 1752598 | 947716 | 15897 | 56 | 3.36 | 0.2515 |

| | | | | | | | | | | |
|---|---|---|---|---|---|---|---|---|---|---|
| LL071_1 | TGACCAAT | GGAACGA | Shortleaf | 2996027 | 2990865 | 1609736 | 20659 | 71 | 0.42 | 0.2879 |
| LL071_2R | ATCACGAT | TATTCGCAT | Shortleaf | 2949631 | 2943080 | 1588692 | 23538 | 61 | 0.51 | 0.2286 |
| LL072 | TTAGGCAT | TATGT | Virginia | 971865 | 955411 | 561148 | 15751 | 32 | 5.89 | 0.2036 |
| LL073_1 | GTCCGCAC | GCAAGCCAT | Shortleaf | 837881 | 836543 | 473630 | 15094 | 27 | 5.17 | 0.2582 |
| LL073_2R | CGATGTAT | AAGACGCT | Shortleaf | 643236 | 642248 | 376038 | 11688 | 28 | 23.52 | 0.2379 |
| LL074.1R | AGTCAACA | TCAGAGAT | Shortleaf | 4151725 | 4144886 | 2037682 | 20254 | 92 | 0.36 | 0.2705 |
| LL074_2 | GTGAAACG | CTTGA | Shortleaf | 1849637 | 1815570 | 955424 | 18494 | 45 | 0.31 | 0.2708 |
| LL074_5 | ATCACGAT | ACAACCAACT | Shortleaf | 2121052 | 2117733 | 1183751 | 21261 | 50 | 0.40 | 0.2587 |
| LL078_1 _Redo | TTAGGCAT | TCACGGAAG | Pitch | 401208 | 400581 | 238925 | 10089 | 18 | 35.88 | 0.2643 |
| LL078_2R | TTAGGCAT | TGGCAACAGA | Pitch | 322359 | 321857 | 193512 | 6994 | 18 | 58.65 | 0.2657 |
| LL078_3 | TTAGGCAT | GGTGCACATT | Pitch | 1718658 | 1715369 | 924532 | 18458 | 46 | 0.65 | 0.2693 |
| LL092R_02 | GTGAAACG | AACGCACATT | Virginia | 1113313 | 1111422 | 635873 | 17296 | 33 | 1.47 | 0.2410 |
| LL102 | TTAGGCAT | AACGCACATT | Pitch | 1623457 | 1620440 | 858349 | 18453 | 43 | 0.74 | 0.2749 |
| LL136 | TGACCAAT | TCACGGAAG | Virginia | 95901 | 95677 | 56150 | 663 | 12 | 95.61 | 0.2784 |
| LL137 | CGATGTAT | CTCGCGG | Virginia | 1968589 | 1963700 | 1075283 | 16316 | 62 | 4.17 | 0.2500 |
| LL139 | CGATGTAT | GAGCGACAT | Shortleaf | 58818 | 56184 | 33052 | 382 | 13 | 97.45 | 0.2000 |
| LL140 | GTGAAACG | TCACGGAAG | Virginia | 2697679 | 2693100 | 1464891 | 20809 | 64 | 0.30 | 0.2693 |
| LL179 | TTAGGCAT | GGAACGA | Shortleaf | 106355 | 106138 | 62382 | 1015 | 13 | 93.37 | 0.2481 |
| RB076_01 | GTGAAACG | TGACGCCA | Virginia | 1121727 | 1120034 | 626912 | 17470 | 32 | 0.81 | 0.2897 |
| RB122_01 ♂ | TTAGGCAT | TAGCGGAT | Shortleaf | 1704230 | 1701472 | 912234 | 17471 | 48 | 2.69 | 0.0035 |
| RB126_01 | GTGAAACG | AACGTGCCT | Virginia | 1383786 | 1381717 | 791328 | 17901 | 40 | 0.55 | 0.2101 |
| RB127_01 | GTCCGCAC | CAGATA | Virginia | 2262967 | 2241692 | 1212993 | 18612 | 58 | 0.52 | 0.2427 |
| RB128_01 | TTAGGCAT | GGCTTA | Virginia | 737594 | 730143 | 404386 | 14891 | 24 | 8.76 | 0.2250 |
| RB129_01 | TTAGGCAT | TCAGAGAT | Shortleaf | 3639318 | 3633285 | 1878860 | 19951 | 88 | 0.29 | 0.2433 |
| RB132C | ATCACGAT | CGTCGCCACT | Pitch | 1978013 | 1975427 | 1085117 | 20112 | 48 | 0.59 | 0.2524 |
| RB135b | CGATGTAT | GGTGCACATT | Virginia | 21960 | 21922 | 13146 | 5 | 36 | 100 | N/A |
| RB337 | TTAGGCAT | TCTTGG | Virginia | 942619 | 929301 | 532282 | 15500 | 30 | 7.89 | 0.1930 |
| RB339 | GTCCGCAC | CCTTGCCATT | Shortleaf | 2050440 | 2048052 | 1123558 | 18264 | 56 | 0.81 | 0.2756 |
| RB343_01 | ATCACGAT | GGAACGA | Pitch | 2370469 | 2366356 | 1295346 | 21518 | 54 | 0.37 | 0.2686 |
| RB344 | AGTCAACA | CCTTGCCATT | Pitch | 2552531 | 2549248 | 1338625 | 18807 | 65 | 0.56 | 0.2208 |
| RB347 | CGATGTAT | ATAGAT | Virginia | 588823 | 582151 | 336710 | 11106 | 26 | 27.87 | 0.1869 |

| | | | | | | | | | | |
|---|---|---|---|---|---|---|---|---|---|---|
| RB349 | GTGAAACG | ATTAT | Shortleaf | 124338 | 121564 | 72081 | 496 | 11 | 96.66 | 0.2425 |
| RB353 | AGTCAACA | TATGT | Shortleaf | 14380 | 11910 | 7239 | 1 | 28 | 100 | N/A |
| RB355 | AGTCAACA | GGTGT | Virginia | 5875532 | 5779433 | 2694037 | 22298 | 109 | 0.36 | 0.2516 |
| RB357 | GTCCGCAC | CCACTCA | Virginia | 1416972 | 1410985 | 761499 | 16976 | 40 | 0.90 | 0.2570 |
| RB358 | TTAGGCAT | CTTGA | Virginia | 987908 | 953466 | 524248 | 16435 | 29 | 3.21 | 0.2575 |

**Table S4. Sequences for ddRAD adapters.** The variable-length barcode incorporated into the P1 barcode is given in the name. The P2 barcode is biotinylated.

| Name | Sequence |
|---|---|
| P1.1_ATTAT | ACACTCTTTCCCTACACGACGCTCTTCCGATCTATTATCATG |
| P1.1_CACCA | ACACTCTTTCCCTACACGACGCTCTTCCGATCTCACCACATG |
| P1.1_CCTCG | ACACTCTTTCCCTACACGACGCTCTTCCGATCTCCTCGCATG |
| P1.1_CTTGA | ACACTCTTTCCCTACACGACGCTCTTCCGATCTCTTGACATG |
| P1.1_GGATA | ACACTCTTTCCCTACACGACGCTCTTCCGATCTGGATACATG |
| P1.1_GGTGT | ACACTCTTTCCCTACACGACGCTCTTCCGATCTGGTGTCATG |
| P1.1_TATGT | ACACTCTTTCCCTACACGACGCTCTTCCGATCTTATGTCATG |
| P1.1_TGCTT | ACACTCTTTCCCTACACGACGCTCTTCCGATCTTGCTTCATG |
| P1.1_AACTGG | ACACTCTTTCCCTACACGACGCTCTTCCGATCTAACTGGCATG |
| P1.1_ACAACT | ACACTCTTTCCCTACACGACGCTCTTCCGATCTACAACTCATG |
| P1.1_ATAGAT | ACACTCTTTCCCTACACGACGCTCTTCCGATCTATAGATCATG |
| P1.1_CAGATA | ACACTCTTTCCCTACACGACGCTCTTCCGATCTCAGATACATG |
| P1.1_GAAGTG | ACACTCTTTCCCTACACGACGCTCTTCCGATCTGAAGTGCATG |
| P1.1_GGCTTA | ACACTCTTTCCCTACACGACGCTCTTCCGATCTGGCTTACATG |
| P1.1_TCTTGG | ACACTCTTTCCCTACACGACGCTCTTCCGATCTTCTTGGCATG |
| P1.1_TCACTG | ACACTCTTTCCCTACACGACGCTCTTCCGATCTTCACTGCATG |
| P1.1_ACCAGGA | ACACTCTTTCCCTACACGACGCTCTTCCGATCTACCAGGACATG |
| P1.1_CCACTCA | ACACTCTTTCCCTACACGACGCTCTTCCGATCTCCACTCACATG |
| P1.1_CCGAACA | ACACTCTTTCCCTACACGACGCTCTTCCGATCTCCGAACACATG |
| P1.1_CTAAGCA | ACACTCTTTCCCTACACGACGCTCTTCCGATCTCTAAGCACATG |
| P1.1_CTCGCGG | ACACTCTTTCCCTACACGACGCTCTTCCGATCTCTCGCGGCATG |
| P1.1_GCGTCCT | ACACTCTTTCCCTACACGACGCTCTTCCGATCTGCGTCCTCATG |
| P1.1_GGAACGA | ACACTCTTTCCCTACACGACGCTCTTCCGATCTGGAACGACATG |
| P1.1_TAGCCAA | ACACTCTTTCCCTACACGACGCTCTTCCGATCTTAGCCAACATG |
| P1.1_ACTGCGAT | ACACTCTTTCCCTACACGACGCTCTTCCGATCTACTGCGATCATG |
| P1.1_ATGAGCAA | ACACTCTTTCCCTACACGACGCTCTTCCGATCTATGAGCAACATG |
| P1.1_GCCTACCT | ACACTCTTTCCCTACACGACGCTCTTCCGATCTGCCTACCTCATG |
| P1.1_TAGCGGAT | ACACTCTTTCCCTACACGACGCTCTTCCGATCTTAGCGGATCATG |
| P1.1_TGACGCCA | ACACTCTTTCCCTACACGACGCTCTTCCGATCTTGACGCCACATG |
| P1.1_ACGGTACT | ACACTCTTTCCCTACACGACGCTCTTCCGATCTACGGTACTCATG |
| P1.1_AAGACGCT | ACACTCTTTCCCTACACGACGCTCTTCCGATCTAAGACGCTCATG |
| P1.1_TCAGAGAT | ACACTCTTTCCCTACACGACGCTCTTCCGATCTTCAGAGATCATG |
| P1.1_ATATCGCCA | ACACTCTTTCCCTACACGACGCTCTTCCGATCTATATCGCCACATG |
| P1.1_GAGCGACAT | ACACTCTTTCCCTACACGACGCTCTTCCGATCTGAGCGACATCATG |
| P1.1_GCAAGCCAT | ACACTCTTTCCCTACACGACGCTCTTCCGATCTGCAAGCCATCATG |
| P1.1_AACGTGCCT | ACACTCTTTCCCTACACGACGCTCTTCCGATCTAACGTGCCTCATG |
| P1.1_TATTCGCAT | ACACTCTTTCCCTACACGACGCTCTTCCGATCTTATTCGCATCATG |
| P1.1_TCACGGAAG | ACACTCTTTCCCTACACGACGCTCTTCCGATCTTCACGGAAGCATG |
| P1.1_TGGCACAGA | ACACTCTTTCCCTACACGACGCTCTTCCGATCTTGGCACAGACATG |
| P1.1_CTCTCGCAT | ACACTCTTTCCCTACACGACGCTCTTCCGATCTCTCTCGCATCATG |

| | |
|---|---|
| P1.1_AACGCACATT | ACACTCTTTCCCTACACGACGCTCTTCCGATCTAACGCACATTCATG |
| P1.1_CCTTGCCATT | ACACTCTTTCCCTACACGACGCTCTTCCGATCTCCTTGCCATTCATG |
| P1.1_CGTCGCCACT | ACACTCTTTCCCTACACGACGCTCTTCCGATCTCGTCGCCACTCATG |
| P1.1_CGTGGACAGT | ACACTCTTTCCCTACACGACGCTCTTCCGATCTCGTGGACAGTCATG |
| P1.1_GGTGCACATT | ACACTCTTTCCCTACACGACGCTCTTCCGATCTGGTGCACATTCATG |
| P1.1_TGGCAACAGA | ACACTCTTTCCCTACACGACGCTCTTCCGATCTTGGCAACAGACATG |
| P1.1_ACAACCAACT | ACACTCTTTCCCTACACGACGCTCTTCCGATCTACAACCAACTCATG |
| P1.1_CAACCACACA | ACACTCTTTCCCTACACGACGCTCTTCCGATCTCAACCACACACATG |
| P1.2_ATTAT | /5Phos/ATAATAGATCGGAAGAGCGTCGTGTAGGGAAAGAGTGT |
| P1.2_CACCA | /5Phos/TGGTGAGATCGGAAGAGCGTCGTGTAGGGAAAGAGTGT |
| P1.2_CCTCG | /5Phos/CGAGGAGATCGGAAGAGCGTCGTGTAGGGAAAGAGTGT |
| P1.2_CTTGA | /5Phos/TCAAGAGATCGGAAGAGCGTCGTGTAGGGAAAGAGTGT |
| P1.2_GGATA | /5Phos/TATCCAGATCGGAAGAGCGTCGTGTAGGGAAAGAGTGT |
| P1.2_GGTGT | /5Phos/ACACCAGATCGGAAGAGCGTCGTGTAGGGAAAGAGTGT |
| P1.2_TATGT | /5Phos/ACATAAGATCGGAAGAGCGTCGTGTAGGGAAAGAGTGT |
| P1.2_TGCTT | /5Phos/AAGCAAGATCGGAAGAGCGTCGTGTAGGGAAAGAGTGT |
| P1.2_AACTGG | /5Phos/CCAGTTAGATCGGAAGAGCGTCGTGTAGGGAAAGAGTGT |
| P1.2_ACAACT | /5Phos/AGTTGTAGATCGGAAGAGCGTCGTGTAGGGAAAGAGTGT |
| P1.2_ATAGAT | /5Phos/ATCTATAGATCGGAAGAGCGTCGTGTAGGGAAAGAGTGT |
| P1.2_CAGATA | /5Phos/TATCTGAGATCGGAAGAGCGTCGTGTAGGGAAAGAGTGT |
| P1.2_GAAGTG | /5Phos/CACTTCAGATCGGAAGAGCGTCGTGTAGGGAAAGAGTGT |
| P1.2_GGCTTA | /5Phos/TAAGCCAGATCGGAAGAGCGTCGTGTAGGGAAAGAGTGT |
| P1.2_TCTTGG | /5Phos/CCAAGAAGATCGGAAGAGCGTCGTGTAGGGAAAGAGTGT |
| P1.2_TCACTG | /5Phos/CAGTGAAGATCGGAAGAGCGTCGTGTAGGGAAAGAGTGT |
| P1.2_ACCAGGA | /5Phos/TCCTGGTAGATCGGAAGAGCGTCGTGTAGGGAAAGAGTGT |
| P1.2_CCACTCA | /5Phos/TGAGTGGAGATCGGAAGAGCGTCGTGTAGGGAAAGAGTGT |
| P1.2_CCGAACA | /5Phos/TGTTCGGAGATCGGAAGAGCGTCGTGTAGGGAAAGAGTGT |
| P1.2_CTAAGCA | /5Phos/TGCTTAGAGATCGGAAGAGCGTCGTGTAGGGAAAGAGTGT |
| P1.2_CTCGCGG | /5Phos/CCGCGAGAGATCGGAAGAGCGTCGTGTAGGGAAAGAGTGT |
| P1.2_GCGTCCT | /5Phos/AGGACGCAGATCGGAAGAGCGTCGTGTAGGGAAAGAGTGT |
| P1.2_GGAACGA | /5Phos/TCGTTCCAGATCGGAAGAGCGTCGTGTAGGGAAAGAGTGT |
| P1.2_TAGCCAA | /5Phos/TTGGCTAAGATCGGAAGAGCGTCGTGTAGGGAAAGAGTGT |
| P1.2_ACTGCGAT | /5Phos/ATCGCAGTAGATCGGAAGAGCGTCGTGTAGGGAAAGAGTGT |
| P1.2_ATGAGCAA | /5Phos/TTGCTCATAGATCGGAAGAGCGTCGTGTAGGGAAAGAGTGT |
| P1.2_GCCTACCT | /5Phos/AGGTAGGCAGATCGGAAGAGCGTCGTGTAGGGAAAGAGTGT |
| P1.2_TAGCGGAT | /5Phos/ATCCGCTAAGATCGGAAGAGCGTCGTGTAGGGAAAGAGTGT |
| P1.2_TGACGCCA | /5Phos/TGGCGTCAAGATCGGAAGAGCGTCGTGTAGGGAAAGAGTGT |
| P1.2_ACGGTACT | /5Phos/AGTACCGTAGATCGGAAGAGCGTCGTGTAGGGAAAGAGTGT |
| P1.2_AAGACGCT | /5Phos/AGCGTCTTAGATCGGAAGAGCGTCGTGTAGGGAAAGAGTGT |
| P1.2_TCAGAGAT | /5Phos/ATCTCTGAAGATCGGAAGAGCGTCGTGTAGGGAAAGAGTGT |
| P1.2_ATATCGCCA | /5Phos/TGGCGATATAGATCGGAAGAGCGTCGTGTAGGGAAAGAGTGT |
| P1.2_GAGCGACAT | /5Phos/ATGTCGCTCAGATCGGAAGAGCGTCGTGTAGGGAAAGAGTGT |
| P1.2_GCAAGCCAT | /5Phos/ATGGCTTGCAGATCGGAAGAGCGTCGTGTAGGGAAAGAGTGT |
| P1.2_AACGTGCCT | /5Phos/AGGCACGTTAGATCGGAAGAGCGTCGTGTAGGGAAAGAGTGT |

| | |
|---|---|
| P1.2_TATTCGCAT | /5Phos/ATGCGAATAAGATCGGAAGAGCGTCGTGTAGGGAAAGAGTGT |
| P1.2_TCACGGAAG | /5Phos/CTTCCGTGAAGATCGGAAGAGCGTCGTGTAGGGAAAGAGTGT |
| P1.2_TGGCACAGA | /5Phos/TCTGTGCCAAGATCGGAAGAGCGTCGTGTAGGGAAAGAGTGT |
| P1.2_CTCTCGCAT | /5Phos/ATGCGAGAGAGATCGGAAGAGCGTCGTGTAGGGAAAGAGTGT |
| P1.2_AACGCACATT | /5Phos/AATGTGCGTTAGATCGGAAGAGCGTCGTGTAGGGAAAGAGTGT |
| P1.2_CCTTGCCATT | /5Phos/AATGGCAAGGAGATCGGAAGAGCGTCGTGTAGGGAAAGAGTGT |
| P1.2_CGTCGCCACT | /5Phos/AGTGGCGACGAGATCGGAAGAGCGTCGTGTAGGGAAAGAGTGT |
| P1.2_CGTGGACAGT | /5Phos/ACTGTCCACGAGATCGGAAGAGCGTCGTGTAGGGAAAGAGTGT |
| P1.2_GGTGCACATT | /5Phos/AATGTGCACCAGATCGGAAGAGCGTCGTGTAGGGAAAGAGTGT |
| P1.2_TGGCAACAGA | /5Phos/TCTGTTGCCAAGATCGGAAGAGCGTCGTGTAGGGAAAGAGTGT |
| P1.2_ACAACCAACT | /5Phos/AGTTGGTTGTAGATCGGAAGAGCGTCGTGTAGGGAAAGAGTGT |
| P1.2_CAACCACACA | /5Phos/TGTGTGGTTGAGATCGGAAGAGCGTCGTGTAGGGAAAGAGTGT |
| P2.1 | GTGACTGGAGTTCAGACGTGTGCTCTTCCGATCT |
| P2.2_biotinylated | /5Phos/AATTAGATCGGAAGAGCGAGAACAA/3Bio/ |

**Table S5. PCR primers for amplification of ddRAD libraries.** The PCR2 primers include standard Illumina multiplexing indexes (given as Index_XXXXXXXX), and also include a 4-base pair string of degenerate bases (N) to allow for PCR duplicate detection. When ordering, the degenerate bases should be requested as "hand mixed" in an equal ratio to prevent over-representation of bases [e.g., replace each "N" when ordering with "(25252525)"].

| Name | Sequence |
|---|---|
| PCR1 | AATGATACGGCGACCACCGAGATCTACACTCTTTCCCTACACGACG |
| PCR2_Index_ATCGTGAT | CAAGCAGAAGACGGCATACGAGATNNNNATCGTGATGTGACTGGAGTTCAGACGTGTGC |
| PCR2_Index_ATACATCG | CAAGCAGAAGACGGCATACGAGATNNNNATACATCGGTGACTGGAGTTCAGACGTGTGC |
| PCR2_Index_ATGCCTAA | CAAGCAGAAGACGGCATACGAGATNNNNATGCCTAAGTGACTGGAGTTCAGACGTGTGC |
| PCR2_Index_ATTGGTCA | CAAGCAGAAGACGGCATACGAGATNNNNATTGGTCAGTGACTGGAGTTCAGACGTGTGC |
| PCR2_Index_ATCACTGT | CAAGCAGAAGACGGCATACGAGATNNNNATCACTGTGTGACTGGAGTTCAGACGTGTGC |
| PCR2_Index_ATGTAGCC | CAAGCAGAAGACGGCATACGAGATNNNNATGTAGCCGTGACTGGAGTTCAGACGTGTGC |
| PCR2_Index_TGTTGACT | CAAGCAGAAGACGGCATACGAGATNNNNTGTTGACTGTGACTGGAGTTCAGACGTGTGC |
| PCR2_Index_CGGGACGG | CAAGCAGAAGACGGCATACGAGATNNNNCGGGACGGGTGACTGGAGTTCAGACGTGTGC |
| PCR2_Index_GTGCGGAC | CAAGCAGAAGACGGCATACGAGATNNNNGTGCGGACGTGACTGGAGTTCAGACGTGTGC |
| PCR2_Index_CGTTTCAC | CAAGCAGAAGACGGCATACGAGATNNNNCGTTTCACGTGACTGGAGTTCAGACGTGTGC |
| PCR2_Index_AAGGCCAC | CAAGCAGAAGACGGCATACGAGATNNNNAAGGCCACGTGACTGGAGTTCAGACGTGTGC |
| PCR2_Index_TCCGAAAC | CAAGCAGAAGACGGCATACGAGATNNNNTCCGAAACGTGACTGGAGTTCAGACGTGTGC |

**Table S6. Statistical results for needle width comparisons.** A. ANOVA table for comparison of needle width as a function of host tree species and age, controlling for individual tree variation. B. Post hoc comparison of differences between host tree species. C. Post hoc comparisons between the mature trees and seedlings of each host. D. Post hoc comparisons between the host species within each host range (mature compared to mature; seedlings compared to seedlings).

**A. Type III ANOVA table for linear mixed model: needle width ~ host species * host age + (1|tree)**

| | SS | MSS | NumDF | DenDF | F | p-value |
|---|---|---|---|---|---|---|
| **host species** | 1.40807 | 0.70404 | 2 | 33 | 87.156 | 6.78E-14 |
| **host age** | 0.78617 | 0.78617 | 1 | 33 | 97.324 | 2.27E-11 |
| **species:age** | 0.84771 | 0.42385 | 2 | 33 | 52.471 | 5.63E-11 |

**B. Post hoc pairwise comparison of host species with FDR correction**

| Contrast | estimate | SE | df | t.ratio | p-value |
|---|---|---|---|---|---|
| rigida – echinata | 0.533 | 0.0407 | 33 | 13.081 | <.0001 |
| rigida - virginiana | 0.397 | 0.0425 | 33 | 9.345 | <.0001 |
| echinata - virgniana | -0.136 | 0.037 | 33 | -3.673 | 0.0008 |

**C. Post hoc comparisons of host age within each host species, with FDR correction**

| species | contrast | estimate | SE | df | t.ratio | p-value |
|---|---|---|---|---|---|---|
| rigida | MatureVsSeedling | 0.8217 | 0.0647 | 33 | 12.698 | <.0001 |
| echinata | MatureVsSeedling | 0.0174 | 0.0494 | 33 | 0.351 | 0.7278 |
| virginiana | MatureVsSeedling | 0.1303 | 0.055 | 33 | 2.369 | 0.0238 |

**D. Post hoc comparisons of host species within each host age, with FDR correction**

| age | contrast | estimate | SE | df | t.ratio | p-value |
|---|---|---|---|---|---|---|
| Mature | rigida - echinata | 0.9347 | 0.0723 | 33 | 12.92 | <.0001 |
| Mature | rigida - virginiana | 0.7425 | 0.0763 | 33 | 9.736 | <.0001 |
| Mature | echinata - virgniana | -0.1923 | 0.0638 | 33 | -3.013 | 0.0049 |
| Seedling | rigida - echinata | 0.1304 | 0.0374 | 33 | 3.49 | 0.0042 |
| Seedling | rigida - virginiana | 0.0511 | 0.0374 | 33 | 1.368 | 0.1806 |
| Seedling | echinata - virgniana | -0.0793 | 0.0374 | 33 | -2.123 | 0.0621 |

**Table S7. Statistical results for resin content comparisons.** A. ANOVA table for comparison of resin content as a function of host tree species and collection month. B. Post hoc comparison of differences between host tree species. C. Post hoc comparisons between the host tree species within each collection month.

**Type III ANOVA table for linear model: sqrt(resin)~host species * collection month**

| | **Sum Sq** | **DF** | **F** | **p-value** |
|---|---|---|---|---|
| **(Intercept)** | 111.48 | 1 | 606.988 | < 2.2e-16 |
| Host | 6.355 | 2 | 17.302 | 1.13E-07 |
| Month | 5.372 | 1 | 29.247 | 1.76E-07 |
| Host:Month | 5.37 | 2 | 14.62 | 1.16E-06 |
| Residuals | 37.834 | 206 | | |

**Post-hoc pairwise comparison of host species with FDR correction**

| **Contrast** | **estimate** | **SE** | **df** | **t.ratio** | **p-value** |
|---|---|---|---|---|---|
| rigida - echinata | 0.21662 | 0.0742 | 206 | 2.92 | 0.009 |
| rigida - virginiana | 0.00989 | 0.0694 | 206 | 0.143 | 0.8868 |
| echinata - virgniana | -0.20673 | 0.0744 | 206 | -2.778 | 0.009 |

**Post-hoc pairwise comparison of host species within sampling months with FDR correction**

| **month** | **contrast** | **estimate** | **SE** | **df** | **t.ratio** | **p.value** |
|---|---|---|---|---|---|---|
| May | rigida - echinata | 0.471 | 0.1063 | 206 | 4.432 | <.0001 |
| May | rigida - virginiana | -0.1365 | 0.1003 | 206 | -1.361 | 0.175 |
| May | echinata - virgniana | -0.6076 | 0.1069 | 206 | -5.682 | <.0001 |
| August | rigida - echinata | -0.0378 | 0.1035 | 206 | -0.365 | 0.7155 |
| August | rigida - virginiana | 0.1563 | 0.0958 | 206 | 1.631 | 0.1566 |
| August | echinata - virgniana | 0.1941 | 0.1035 | 206 | 1.875 | 0.1566 |

**Table S8. Statistical results for ovipositor morphology analysis.** A. Proportion of variation explained by each PC of ovipositor shape. B. Procrustes AMOVA table for landmark coordinates as a function of source host. C. Post hoc comparisons between source hosts.

**A. Importance of components**

| | **Comp1** | **Comp2** | **Comp3** | **Comp4** | **Comp5** | **Comp6** | **Comp7** |
|---|---|---|---|---|---|---|---|
| **Eigenvalues** | 0.00020058 | 8.44E-05 | 6.95E-05 | 5.99E-05 | 4.89E-05 | 4.35E-05 | 3.33E-05 |
| **Proportion of variance** | 0.27440435 | 1.15E-01 | 9.50E-02 | 8.19E-02 | 6.68E-02 | 5.95E-02 | 4.55E-02 |
| **Cumulative Proportion** | 0.27440435 | 3.90E-01 | 4.85E-01 | 5.67E-01 | 6.34E-01 | 6.93E-01 | 7.39E-01 |
| | **Comp8** | **Comp9** | **Comp10** | **Comp11** | **Comp12** | **Comp13** | **Comp14** |
| **Eigenvalues** | 2.77E-05 | 2.37E-05 | 2.05E-05 | 1.84E-05 | 1.53E-05 | 1.34E-05 | 1.1087E-05 |
| **Proportion of variance** | 3.79E-02 | 3.25E-02 | 2.80E-02 | 2.51E-02 | 2.09E-02 | 1.83E-02 | 0.01516771 |
| **Cumulative Proportion** | 7.77E-01 | 8.09E-01 | 8.37E-01 | 8.62E-01 | 8.83E-01 | 9.01E-01 | 0.91660311 |
| | **Comp15** | **Comp16** | **Comp17** | **Comp18** | **Comp19** | **Comp20** | **Comp21** |
| **Eigenvalues** | 9.78E-06 | 8.09E-06 | 7.77E-06 | 7.15E-06 | 5.77E-06 | 5.11E-06 | 4.71E-06 |
| **Proportion of variance** | 1.34E-02 | 1.11E-02 | 1.06E-02 | 9.78E-03 | 7.90E-03 | 6.99E-03 | 6.44E-03 |
| **Cumulative Proportion** | 9.30E-01 | 9.41E-01 | 9.52E-01 | 9.61E-01 | 9.69E-01 | 9.76E-01 | 9.83E-01 |
| | **Comp22** | **Comp23** | **Comp24** | **Comp25** | **Comp26** | **Comp27** | |
| **Eigenvalues** | 3.02E-06 | 2.83E-06 | 2.43E-06 | 2.06E-06 | 1.20E-06 | 1.06E-06 | |
| **Proportion of variance** | 4.13E-03 | 3.87E-03 | 3.32E-03 | 2.81E-03 | 1.64E-03 | 1.44E-03 | |
| **Cumulative Proportion** | 9.87E-01 | 9.91E-01 | 9.94E-01 | 9.97E-01 | 9.99E-01 | 1.00E+00 | |

**B. Procrustes ANOVA (MANOVA) Type I (Sequential) Sums of Squares and Cross-products (landmark_coordinates ~ host species)**

| | **df** | **SS** | **MS** | **Rsq** | **F** | **P.val** |
|---|---|---|---|---|---|---|
| **Host** | 2 | 0.0034626 | 0.00173128 | 0.17544 | 2.6596 | 0.00059994 |
| **Residuals** | 25 | 0.0162737 | 0.00065095 | | | |
| **Total** | 27 | 0.0197363 | 0.00073097 | | | |

**C. Post hoc pairwise D comparison for host species**

| | df | SS | MS | Rsq | F | P.val |
|---|---|---|---|---|---|---|
| Host | 2 | 0.0034626 | 0.00173128 | 0.17544 | 2.6596 | 0.00049995 |
| Residuals | 25 | 0.0162737 | 0.00065095 | | | |
| Total | 27 | 0.0197363 | 0.00073097 | | | |

| Observed distance | Pitch | Shortleaf | Virginia |
|---|---|---|---|
| Pitch | 0 | 0.02319688 | 0.01175813 |
| Shortleaf | 0.02319688 | 0 | 0.0200255 |
| Virginia | 0.01175813 | 0.0200255 | 0 |

| P-value | Pitch | Shortleaf | Virginia |
|---|---|---|---|
| Pitch | 1 | 0.00029997 | 0.6030397 |
| Shortleaf | 0.00029997 | 1 | 0.00819918 |
| Virginia | 0.6030397 | 0.00819918 | 1 |

**Table S9. Statistical results for ovipositor length and width comparisons.** ANOVA tables for ovipositor length (A) and width (B) as a function of source host.

**A. Type II ANOVA for ovipositor length ~ host species**

| | **Sum Sq** | **Df** | **F value** | **Pr(>F)** |
|---|---|---|---|---|
| Host | 0.000562 | 2 | 0.7057 | 0.5033 |
| Residuals | 0.009949 | 25 | | |

**B. Type II ANOVA for ovipositor width ~ host species**

| | **Sum Sq** | **Df** | **F value** | **Pr(>F)** |
|---|---|---|---|---|
| Host | 0.000167 | 2 | 0.9925 | 0.3848 |
| Residuals | 0.002107 | 25 | | |

**Table S10. Statistical results for larval survival comparisons.** A. ANOVA table for larval survival to cocoon stage as a function of sawfly line and rearing host, controlling for colony level variation. B. Post hoc pairwise comparisons of larval survival between sawfly lines. C. Post hoc pairwise comparisons of larval survival on different rearing hosts within sawfly lines.

**A. Type III ANOVA table for generalized linear mixed model: Survival~Line*Host+(1|ColonyID), family=binomial**

| | **Chisq** | **DF** | **p-value** |
|---|---|---|---|
| **(Intercept)** | 8.0195 | 1 | 0.004628 |
| Line | 12.2965 | 2 | 0.002137 |
| Host | 4.5509 | 2 | 0.102752 |
| Line:Host | 9.6459 | 4 | 0.046835 |

**B. Post-hoc pairwise comparison of sawfly line with FDR correction**

| **contrast** | **estimate** | **SE** | **DF** | **z.ratio** | **p.value** |
|---|---|---|---|---|---|
| Pitch-Shortleaf | -1.055 | 0.384 | Inf | -2.746 | 0.0181 |
| Pitch-Virginia | -0.315 | 0.364 | Inf | -0.866 | 0.3867 |
| Shortleaf-Virginia | 0.74 | 0.383 | Inf | 1.93 | 0.0804 |

**C. Post-hoc pairwise comparison of rearing hosts within sawfly lines with FDR correction**

| **Line** | **contrast** | **estimate** | **SE** | **DF** | **z.ratio** | **p.value** |
|---|---|---|---|---|---|---|
| Pitch | rigida - echinata | -1.3016 | 0.622 | Inf | -2.092 | 0.1092 |
| Pitch | rigida - virginiana | -0.6543 | 0.689 | Inf | -0.95 | 0.342 |
| Pitch | echinata - virginiana | 0.6472 | 0.581 | Inf | 1.114 | 0.342 |
| Shortleaf | rigida - echinata | -0.1662 | 0.696 | Inf | -0.239 | 0.8113 |
| Shortleaf | rigida - virginiana | 0.2171 | 0.698 | Inf | 0.311 | 0.8113 |
| Shortleaf | echinata - virginiana | 0.3832 | 0.697 | Inf | 0.55 | 0.8113 |
| Virginia | rigida - echinata | -2.4897 | 0.664 | Inf | -3.749 | 0.0003 |
| Virginia | rigida - virginiana | -2.4742 | 0.647 | Inf | -3.827 | 0.0003 |
| Virginia | echinata - virginiana | 0.0155 | 0.574 | Inf | 0.027 | 0.9785 |

**Table S11. Statistical results for development time comparisons.** A. ANOVA table for development time as a function of sawfly line and rearing host, controlling for colony level variation. B. Post hoc pairwise comparisons of development time between sawfly lines. C. Post hoc pairwise comparisons of development time between rearing hosts. D. Post hoc pairwise comparisons of development time between rearing hosts within sawfly lines.

**A. Type III ANOVA table for generalized linear mixed model:DevTim~Line*Host+(1|ColonyID), family=Gamma**

| | Chisq | Df | p-value |
|---|---|---|---|
| **(Intercept)** | 4990.955 | 1 | < 2.2e-16 |
| Line | 20.707 | 2 | 3.19E-05 |
| Host | 20.407 | 2 | 3.71E-05 |
| Line:Host | 70.764 | 4 | 1.57E-14 |

**B. Post-hoc pairwise comparison of sawfly line with FDR correction**

| contrast | estimate | SE | df | z.ratio | p.value |
|---|---|---|---|---|---|
| Pitch-Shortleaf | -1.02E-03 | 0.000234 | Inf | -4.363 | <.0001 |
| Pitch-Virginia | 2.45E-05 | 0.00023 | Inf | 0.106 | 0.9152 |
| Shortleaf-Virginia | 1.04E-03 | 0.000228 | Inf | 4.578 | <.0001 |

**C. Post-hoc pairwise comparison of rearing host with FDR correction**

| contrast | estimate | SE | df | z.ratio | p.value |
|---|---|---|---|---|---|
| rigida - echinata | -0.001036 | 0.000237 | Inf | -4.374 | <.0001 |
| rigida - virginiana | -0.000479 | 0.000242 | Inf | -1.983 | 0.0474 |
| echinata - virginiana | 0.000557 | 0.000212 | Inf | 2.623 | 0.0131 |

**D. Post-hoc pairwise comparison of rearing host within each sawfly line, with FDR correction**

| Line | contrast | estimate | SE | df | z.ratio | p.value |
|---|---|---|---|---|---|---|
| Pitch | rigida - echinata | 0.001006 | 0.000414 | Inf | 2.428 | 0.0152 |
| Pitch | rigida - virginiana | 0.001984 | 0.000443 | Inf | 4.48 | <.0001 |
| Pitch | echinata - virginiana | 0.000978 | 0.000364 | Inf | 2.687 | 0.0108 |
| Shortleaf | rigida - echinata | -0.002708 | 0.000399 | Inf | -6.794 | <.0001 |
| Shortleaf | rigida - virginiana | -0.001364 | 0.000403 | Inf | -3.387 | 0.0008 |
| Shortleaf | echinata - virginiana | 0.001344 | 0.000403 | Inf | 3.337 | 0.0008 |
| Virginia | rigida - echinata | -0.001407 | 0.000418 | Inf | -3.367 | 0.0011 |
| Virginia | rigida - virginiana | -0.002059 | 0.00041 | Inf | -5.025 | <.0001 |
| Virginia | echinata - virginiana | -0.000651 | 0.000333 | Inf | -1.956 | 0.0505 |

**Table S12. Statistical results for female cocoon weight comparisons.** A. Type III ANOVA table for female cocoon weight as a function of sawfly line and rearing host, controlling for colony level variation. B. Type II ANOVA table for female cocoon weight as a function of sawfly line and rearing host, controlling for colony level variation. C. Post hoc pairwise comparisons of female cocoon weight between sawfly lines. D. Post hoc pairwise comparisons of female cocoon weight between rearing hosts.

**A. Type III ANOVA table for linear mixed model:FemCocoonWeight~Line*Host+(1|ColonyID)**

| | SS | MSS | NumDF | DenDF | F | p-value |
|---|---|---|---|---|---|---|
| Line | 0.00055579 | 2.78E-04 | 2 | 33.852 | 7.2065 | 0.002469 |
| Host | 0.000472 | 2.36E-04 | 2 | 33.884 | 6.12 | 0.005382 |
| Line*Host | 0.0001462 | 3.65E-05 | 4 | 33.832 | 0.9478 | 0.448539 |

**B. Type II ANOVA table for linear mixed model: FemCocoonWeight~Line+Host+(1|ColonyID)**

| | SS | MSS | NumDF | DenDF | F | p-value |
|---|---|---|---|---|---|---|
| Line | 0.00070048 | 0.00035024 | 2 | 37.509 | 9.0814 | 0.0006076 |
| Host | 0.00056137 | 0.00028068 | 2 | 37.405 | 7.2779 | 0.0021394 |

**C. Post-hoc pairwise comparison of sawfly line with FDR correction**

| contrast | estimate | SE | df | t.ratio | p.value |
|---|---|---|---|---|---|
| Pitch-Shortleaf | 0.00572 | 0.00393 | 38 | 1.454 | 0.1541 |
| Pitch-Virginia | -0.00995 | 0.00378 | 37.9 | -2.631 | 0.0184 |
| Shortleaf-Virginia | -0.01567 | 0.00375 | 38.1 | -4.181 | 0.0005 |

**D. Post hoc pairwise comparison of rearing host with FDR correction.**

| contrast | estimate | SE | df | t.ratio | p.value |
|---|---|---|---|---|---|
| rigida - echinata | 0.00917 | 0.004 | 37.7 | 2.293 | 0.0413 |
| rigida - virginiana | -0.00435 | 0.00398 | 37.8 | -1.091 | 0.282 |
| echinata - virginiana | -0.01352 | 0.0036 | 38.2 | -3.758 | 0.0017 |

**Table S13. Statistical results for male cocoon weight comparisons.** A. Type III ANOVA table for male cocoon weight as a function of sawfly line and rearing host, controlling for colony level variation. B. Type II ANOVA table for male cocoon weight as a function of sawfly line and rearing host, controlling for colony level variation. C. Post hoc pairwise comparisons of male cocoon weight between sawfly lines. D. Post hoc pairwise comparisons of male cocoon weight between rearing hosts.

**A. Type III ANOVA table for linear mixed model:MaleCocoonWeight~Line*Host+(1|ColonyID)**

| | SS | MSS | NumDF | DenDF | F | p-value |
|---|---|---|---|---|---|---|
| Line | 0.0004549 | 2.27E-04 | 2 | 30.824 | 18.4295 | 5.44E-06 |
| Host | 0.00025145 | 1.26E-04 | 2 | 31.026 | 10.187 | 0.0003972 |
| Line:Host | 0.00006113 | 1.53E-05 | 4 | 30.71 | 1.2382 | 0.3153537 |

**B. Type II ANOVA table for linear mixed model: FemCocoonWeight~Line+Host+(1|ColonyID)**

| | SS | MSS | NumDF | DenDF | F | p-value |
|---|---|---|---|---|---|---|
| Line | 0.00056988 | 0.00028494 | 2 | 33.661 | 23.058 | 4.93E-07 |
| Host | 0.00027918 | 0.00013959 | 2 | 33.188 | 11.296 | 0.0001812 |

**C. Post-hoc pairwise comparison of sawfly line with FDR correction**

| contrast | estimate | SE | df | t.ratio | p.value |
|---|---|---|---|---|---|
| Pitch-Shortleaf | 0.00566 | 0.00113 | 33.4 | 5.032 | <.0001 |
| Pitch-Virginia | -0.00129 | 0.00113 | 38.2 | -1.134 | 0.2638 |
| Shortleaf-Virginia | -0.00695 | 0.00109 | 33.2 | -6.398 | <.0001 |

**D. Post hoc pairwise comparison of rearing host with FDR correction**

| contrast | estimate | SE | df | t.ratio | p.value |
|---|---|---|---|---|---|
| rigida - echinata | 6.33E-05 | 0.00113 | 32.7 | 0.056 | 0.9556 |
| rigida - virginiana | -4.54E-03 | 0.0012 | 34.4 | -3.773 | 0.0009 |
| echinata - virginiana | -4.61E-03 | 0.00105 | 35.7 | -4.393 | 0.0003 |

**Table S14. Statistical results for mating outcome comparisons.** ANOVA table for mating outcome (yes/no) as a function of female and male line. There was no impact of male or female line on mating, nor was there an interaction between female or male line.

**Type III ANOVA table for generalized linear model: MatingOutcome~FemaleLine*MaleLine, family=binomial**

| | **Chisq** | **Df** | **p-value** |
|---|---|---|---|
| Female | 1.4868 | 2 | 0.4755 |
| Male | 2.6561 | 2 | 0.265 |
| Female:Male | 3.4523 | 4 | 0.4852 |

**Table S15. Statistical results for hybrid survival outcome comparisons.** ANOVA table for hybrid survival outcome (yes/no) as a function of female and male line. There was no impact of male or female line on mating, nor was there an interaction between female or male line.

**Type III ANOVA table for generalized linear model: Hybrid Survival~FemaleLine*MaleLine, family=binomial**

| | **Chisq** | **Df** | **Pr(>Chisq)** |
|---|---|---|---|
| (Intercept) | 0.0002 | 1 | 0.9878 |
| Female | 1.4424 | 2 | 0.4862 |
| Male | 1.7455 | 2 | 0.4178 |
| Female:Male | 6.9489 | 4 | 0.1386 |

**Table S16. Statistical results for no-choice host preference assays.** A. Type III ANOVA table for laying outcome (yes/no) as a function of sawfly line and offered host. B. Type II ANOVA table for laying outcome (yes/no) as a function of sawfly line. C. Post hoc pairwise comparisons of laying outcome between sawfly lines.

**A. Type III ANOVA table for generalized linear model: LayingOutcome~Line*Host, family=binomial**

| | Chisq | Df | p-value |
|---|---|---|---|
| Line | 6.4973 | 2 | 0.03883 |
| Host | 1.5557 | 2 | 0.4594 |
| Line:Host | 8.0252 | 4 | 0.09066 |

**B. Type II ANOVA table for reduced gneralized linear model: LayingOutcome~Line, family=binomial**

| | Chisq | Df | p-value |
|---|---|---|---|
| line | 16.01 | 2 | 0.0003338 |

**C. Post-hoc pairwise comparison of sawfly line with FDR correction**

| contrast | estimate | SE | df | z.ratio | p.value |
|---|---|---|---|---|---|
| Pitch-Shortleaf | -1.255 | 0.401 | Inf | -3.129 | 0.0053 |
| Pitch-Virginia | -0.261 | 0.355 | Inf | -0.735 | 0.4626 |
| Shortleaf-Virginia | 0.994 | 0.415 | Inf | 2.397 | 0.0248 |

**Table S17. Statistical results of choice host preference assays.** Each sawfly line was assessed for deviation from 50:50 host choice (which would be representative of having no preference for their source *vs* non-source hosts) using a separate binomial test.

| **Line** | **# trials** | **# trials with eggs** | **# trails with eggs on source host** | **one-tailed p-value** | **% laid** |
|---|---|---|---|---|---|
| Pitch | 70 | 38 | 14 | 0.9635 | 0.54285714 |
| Shortleaf | 62 | 50 | 27 | 0.3359 | 0.80645161 |
| Virginia | 61 | 37 | 25 | 0.02352 | 0.60655738 |
